# Genomic insights into the host specific adaptation of the *Pneumocystis* genus and emergence of the human pathogen *Pneumocystis jirovecii*

**DOI:** 10.1101/2020.07.29.227421

**Authors:** Ousmane H. Cissé, Liang Ma, John P. Dekker, Pavel P. Khil, Jung-Ho Youn, Jason M. Brenchley, Robert Blair, Bapi Pahar, Magali Chabé, Koen K.A. Van Rompay, Rebekah Keesler, Antti Sukura, Vanessa Hirsch, Geetha Kutty, Yueqin Liu, Peng Li, Jie Chen, Jun Song, Christiane Weissenbacher-Lang, Jie Xu, Nathan S. Upham, Jason E. Stajich, Christina A. Cuomo, Melanie T. Cushion, Joseph A. Kovacs

**Affiliations:** Critical Care Medicine Department, NIH Clinical Center, National Institutes of Health, Bethesda, Maryland, USA; Bacterial Pathogenesis and Antimicrobial Resistance Unit, National Institute of Allergy and Infectious Diseases, National Institutes of Health, Bethesda, Maryland, USA; Department of Laboratory Medicine, NIH Clinical Center, National Institutes of Health, Bethesda, Maryland, USA; Laboratory of Viral Diseases, National Institute of Allergy and Infectious Diseases, National Institutes of Health, Bethesda, Maryland, USA; Tulane National Primate Research Center, Tulane University, New Orleans, Louisiana, USA; Univ. Lille, CNRS, Inserm, CHU Lille, Institut Pasteur de Lille, U1019-UMR 9017-CIIL-Centre d’Infection et d’Immunité de Lille, Lille, France; California National Primate Research Center, University of California, Davis, California, USA; Department of Veterinary Pathology, Faculty of Veterinary Medicine, University of Helsinki, Helsinki, Finland; Laboratory of Molecular Microbiology, National Institute of Allergy and Infectious Disease, National Institutes of Health, Bethesda, Maryland, USA; Department of Respiratory and Critical Care Medicine, the First Affiliated Hospital of Chongqing Medical University, Chongqing, China; Center for Advanced Models for Translational Sciences and Therapeutics, University of Michigan Medical Center, University of Michigan Medical School, Ann Arbor, MI, United States; Institute of Pathology and Forensic Veterinary Medicine, Department of Pathobiology, University of Veterinary Medicine Vienna, Vienna, Austria; Arizona State University, School of Life Sciences, Tempe, Arizona, USA; Department of Microbiology and Plant Pathology and Institute for Integrative Genome Biology, University of California, Riverside, Riverside-California, Riverside, USA; Broad Institute of Harvard and Massachusetts Institute of Technology, Cambridge, Massachusetts, USA; Department of Internal Medicine, College of Medicine, University of Cincinnati, Cincinnati, OH, USA

**Author notes:** These authors contributed equally to this work (Ousmane H. Cissé, Liang Ma). Correspondence and requests for materials should be addressed to O.H.C or L.M or J.A.K.

## Abstract

*Pneumocystis jirovecii*, the fungal agent of human *Pneumocystis* pneumonia, is closely related to macaque *Pneumocystis*. Little is known about other *Pneumocystis* species in distantly related mammals, none of which are capable of establishing infection in humans. The molecular basis of host specificity in *Pneumocystis* remains unknown as experiments are limited due to an inability to culture any species *in vitro*. To explore *Pneumocystis* evolutionary adaptations, we have sequenced the genomes of species infecting macaques, rabbits, dogs and rats and compared them to available genomes of species infecting humans, mice and rats. Complete whole genome sequence data enables analysis and robust phylogeny, identification of important genetic features of the host adaptation, and estimation of speciation timing relative to the rise of their mammalian hosts. Our data reveals novel insights into the evolution of *P. jirovecii*, the sole member of the genus able to infect humans.

## Introduction

The evolutionary history of *Pneumocystis jirovecii*, a fungus that causes life-threatening pneumonia in immunosuppressed patients such as those with HIV infection, has been poorly defined. *P. jirovecii* is derived from a much broader group of host-specific parasites that infect all mammals studied to date. Until recently, *P. carinii* and *P. murina* (which infect rats and mice, respectively) were the only other species in this genus for which biological specimens suitable for whole genome sequencing were readily available. Inter-species inoculation studies have found that a given *Pneumocystis* species can only infect a unique mammalian species (*1, 2*). Further, rats are the only mammals known to be coinfected by at least two distinct *Pneumocystis* species (*P. carinii* and *P. wakefieldiae*) (*3*). Within the *Pneumocystis* genus, *P. jirovecii* is the only species able to infect and reproduce in humans, although the molecular mechanisms of its host adaptation remain elusive.

Previous efforts to reconstruct the evolutionary history of *Pneumocystis* have estimated the origins of the genus at a minimum of 100 million years ago (mya) (*4*). Using a partial transcriptome of *P. macacae*, the *Pneumocystis* species that infects macaques, we recently estimated that *P. jirovecii* diverged from the common ancestor of *P. macacae* around ~62 mya (*5*), which substantially precedes the human-macaque split of ~20 mya ago (*6*). Population bottlenecks in *P. jirovecii* and *P. carinii* at 400,000 and 16,000 years ago, respectively (*5*), are also not concordant with population expansions in modern humans (~200,000 y ago (*7*)) and rats (~10,000 y ago (*8*)), which suggests a decoupled coevolution between *Pneumocystis* and their hosts. This was the first evidence that *Pneumocystis* may not be strictly co-evolving with their mammalian hosts as suggested by ribosomal RNA-based maximum phylogenies (*9*). A molecular clock has not been tested in any of these phylogenies. A strict co-evolution hypothesis was further challenged by evidence showing relaxation of the host specificity in *Pneumocystis* infecting rodents (*10, 11*). However, the accuracy of speciation times is limited without the complete genomes of additional species including that of *P. macacae*, the closest living sister species to *P. jirovecii* identified to date.

The absence of long-term *in vitro* culture methods or animal models for most *Pneumocystis* species has precluded obtaining sufficient DNA for full genome sequencing and has hindered investigation of the *Pneumocystis* genus. So far, only the genomes of human *P. jirovecii* (*12, 13*), rat *P. carinii* (*13, 14*) and mouse *P. murina*, (*13*) are available. These data have provided important insights into the evolution of this genus, including a substantial genome reduction (*12, 13*), the presence of intron-rich genes possibly contributing to transcriptome complexity, and a significant expansion of a highly polymorphic major surface glycoprotein (*msg*) gene superfamily (*13*), some of which are important for immune evasion. However, the lack of whole genome sequences for many species of this genus (particularly the closely related *P. macacae*) has severely constrained the understanding of the implications of these genome features in *Pneumocystis* evolution and adaptation to hosts.

To further explore the evolutionary history of the *Pneumocystis* genus, and explore *P. jirovecii* genetic factors that support its adaptation to humans, we sequenced 2 to 6 specimens of four additional species: those that infect macaques (*P. carinii* forma specialis *macacae* hereafter referred to as *P. macacae*), rabbits (*P. oryctolagi*), dogs (*P. carinii* f. sp. *canis* hereafter referred to as *P. canis*) and rats (*P. wakefieldiae*). We assembled a single representative, nearly full-length genome for three of the species except in *P. canis*, in which we recovered three distinct genome assemblies that appear to represent two separate species. We reconstructed a robust phylogeny of *Pneumocystis* species, estimated their diversification time, and used comparative genomics to identify unique and shared genomic features. Given that wild mammals may be commonly exposed to different *Pneumocystis* species in nature, there is a possibility of historical gene flow among *Pneumocystis* species that we have evaluated.

## Results and Discussion

### Direct sequencing of *Pneumocystis-host* mixed samples

We sequenced the genomes of *Pneumocystis* species from infected macaques, rabbits, dogs and rats (see Methods and Supplementary Methods). Specimens originated from immunosuppressed animals as a consequence of simian immunodeficiency virus infection in macaques, corticosteroid treatment (rabbits and rats), or possible congenital immunodeficiencies (dogs). For each species, we sequenced samples from 2–6 animals (Supplementary Tables 1 and 2). These data were used to assemble one high quality consensus, nearly full-length genome assembly for each species except *P. canis* for which we recovered two nearly full-length assemblies and an additional partial assembly from separate samples (denoted as A, Ck1 and Ck2). Post assembly mapping revealed a non-negligible amount of genetic variability among samples, for example the average genome wide single nucleotide polymorphism (SNP) diversity among six *P. macacae* isolates is ~ 12%. The genome of *P. macacae* was sequenced using Oxford Nanopore long reads and Illumina short read sequences, whereas the other *Pneumocystis* were sequenced only with Illumina (Supplementary Tables 2 and 3). The new *Pneumocystis* genome assemblies range from 7.3 Mb in *P. wakefieldiae* to 8.2 Mb in *P. macacae*. The *P. macacae* and *P. wakefieldiae* genome assemblies consist of 16 and 17 scaffolds, respectively, both of which are highly contiguous and approach the chromosomal level based on similarities with published karyotypes (*3, 15*) and/or the presence of *Pneumocystis* telomere repeats (*16*) at the scaffold ends (Supplementary Table 3). The genome assemblies of *P. oryctolagi* and *P. canis* (assemblies A, Ck1 and Ck2) are less contiguous with 38, 33, 78 and 315 scaffolds, respectively. All these assemblies except for the partial assembly of *P. canis* Ck2 have very similar total sizes (7.3 – 8.2 Mb) comparable to previously sequenced genomes of *P. jirovecii, P. carinii* and *P. murina*, all of which are at or near chromosomal-level with a size of 7.4 – 8.3 Mb (Supplementary Table 3). The genome assemblies are all AT-rich (~71%) and ~3% encodes DNA transposons and retrotransposons (Supplementary Table 3). We also assembled complete mitochondrial genomes from all species in this study, which are similar in size (21.2 – 24.5 kilobases) to published rodent *Pneumocystis* mitogenomes (*24.6 – 26.1 kb*) (*17*) but smaller than that of *P. jirovecii* (~35 kb) (*17*) (Supplementary Table 3). *P. macacae* has a circular mitogenome similar to *P. jirovecii* (*17*) whereas all other sequenced species have linear mitogenomes.

### Genomic differences among *Pneumocystis* species

To assess the extent of genome structure variations among species, we generated whole genome alignment of all representative genome assemblies. We found high levels of interspecies rearrangements ranging from 10 breakpoints between *P. wakefieldiae* and *P. murina* to 142 between *P. jirovecii* and *P. oryctolagi* (Fig. 1; Supplementary Table 4). The vast majority of chromosomal rearrangements were inversions, which, for example accounted for 23 out of 29 breakpoints between *P. jirovecii* and *P. macacae* (Supplementary Table 4). Analysis of aligned raw Nanopore and/or Illumina reads back to the assemblies show no evidence of incorrect contig joins around rearrangement breakpoints. There are clearly less rearrangements among rodent *Pneumocystis* species (*P. wakefieldiae, P. carinii* and *P. murina*) than among all other species (Fig. 1; Supplementary Table 4), which is likely due to their younger evolutionary ages and closer taxonomic relationships of their host species (Figs. 2a and 2c). These rearrangements could have caused incompatibilities between these species, thus preventing gene flow, for species that infect the same host.

**Fig. 1.**
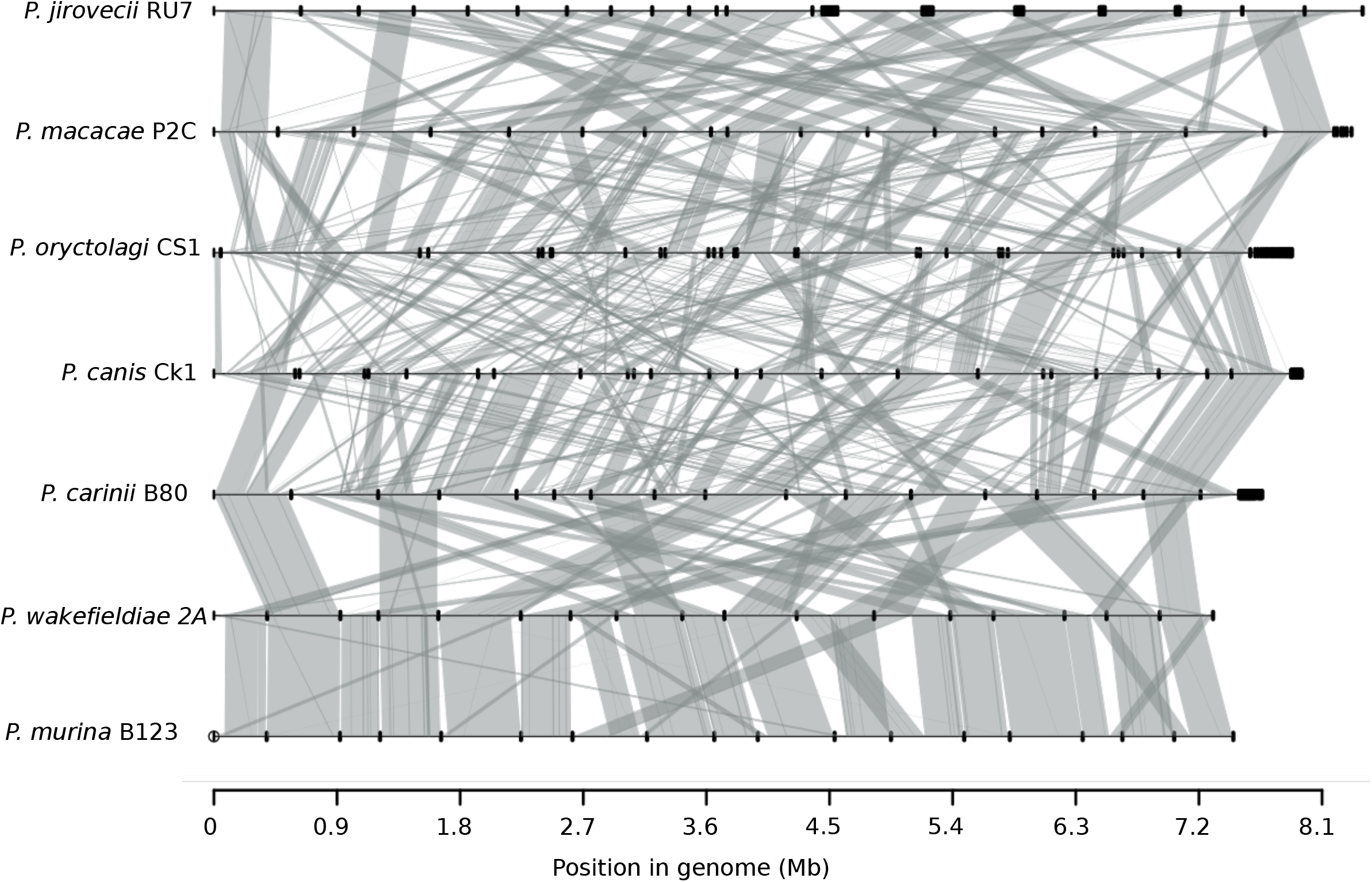
Whole genome structure and synteny among *Pneumocystis* species. Species names and their genome assembly identifiers are shown on the left. Horizontal black lines on the right represent sequences of all scaffolds for each genome laid end-to-end, with their nucleotide positions indicated at the bottom. Dark thick squares represent short scaffolds. Syntenic regions between genomes are linked with vertical gray lines. Reference genome assemblies of *P. jirovecii, P. carinii* and *P. murina* are from a prior study (*13*).

**Fig. 2.**
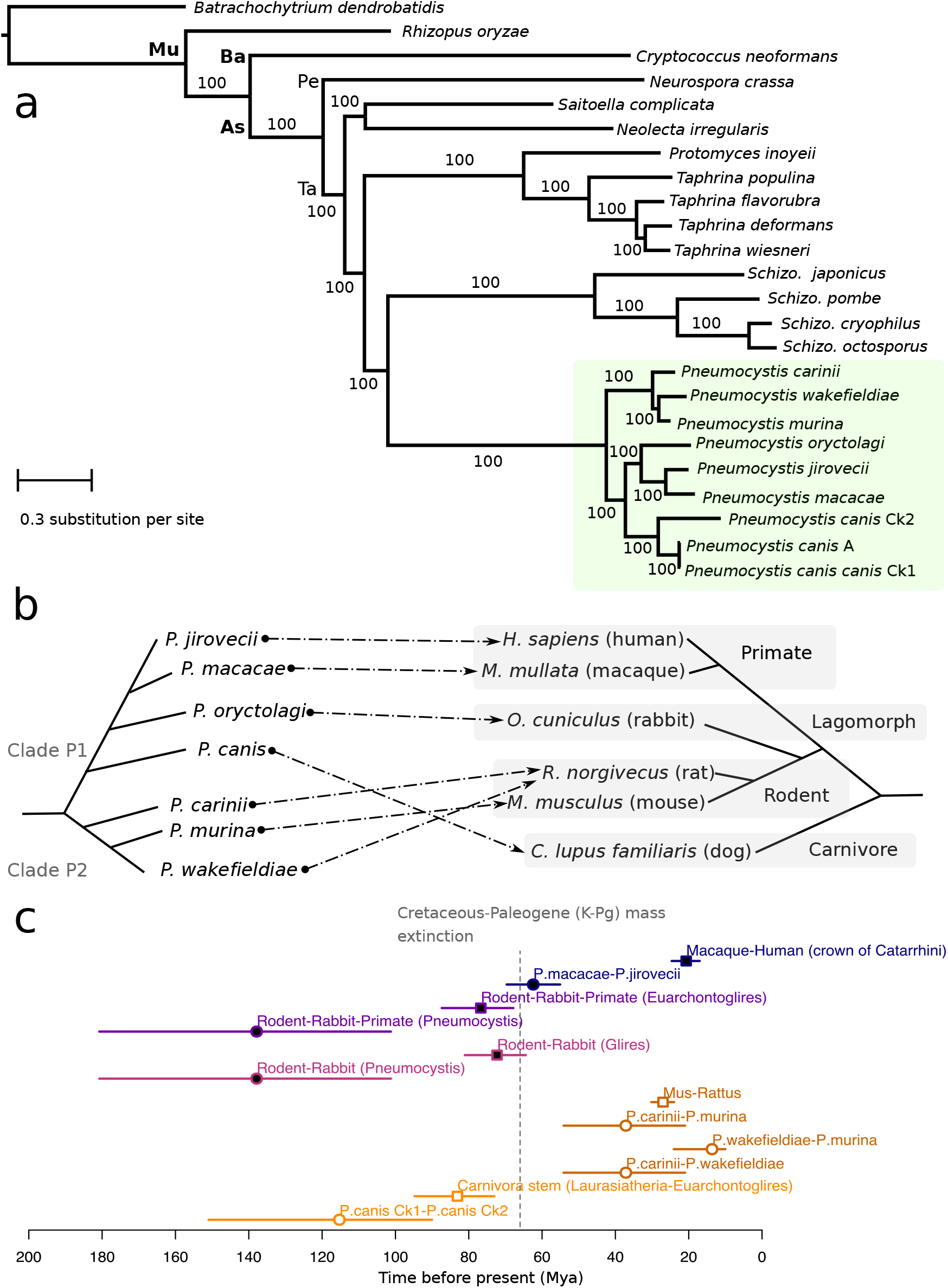
Phylogeny and divergence times of *Pneumocystis* species. a, Maximum likelihood phylogeny constructed using 106 single-copy genes based on 1,000 replicates from 24 annotated fungal genome assemblies including 9 from *Pneumocystis* (highlighted with green background). Only one assembly is shown for each species except there are three for *P. canis* (assemblies Ck1, Ck2 and A). Bootstrap support (%) is presented on the branches. The fungal major phylogenetic phyla and subphyla are represented by their initials: *As* (Ascomycota), *Ba* (Basidiomycota), *Pe* (Pezizomycotina), *Mu* (Mucoromycota) and *Ta* (Taphrinomycotina). b, Schematic representation of species phylogeny and association between *Pneumocystis* species and their respective mammalian hosts. The dashed arrows directed lines represent the specific parasite-host relationships. c, Divergence times of *Pneumocystis* species and mammals. Divergence time medians are represented as squares for hosts and as circles for *Pneumocystis*, and the horizontal lines represent the 95% confidence intervals (CI), which are color-coded the same for each *Pneumocystis* and its host. Closed elements represent nodes that are significantly different in term of divergence times (non-overlapping confidence intervals) whereas open elements represent nodes with overlapping confidence intervals. Catarrhini, taxonomic category (parvorder) including Humans, great apes, gibbons and Old-World monkeys. Euarchontoglires, superorder of mammals including rodents, lagomorphs, treeshrews, colugos and primates. Glires, taxonomic clade consisting of rodents and lagomorphs. Laurasiatheria, taxonomic clade of placental mammals that includes shrews, whales, bats, and carnivorans. Mya, million years ago. K-Pg, Cretaceous-Paleogene. The dotted vertical line representing the K-Pg mass extinction event at 66 mya is included for context only.

Comparison of pairwise whole genome alignment identities between species indicates a substantial genetic divergence: 14% dissimilarity in aligned regions between *P. jirovecii* and *P. macacae;* 21% between *P. jirovecii* and *P. oryctolagi;* 22% between *P. jirovecii* and *P. canis* Ck1; 15% between *P. wakefieldiae* and *P. carinii;* and 12% between *P. wakefieldiae* and *P. murina* (Supplementary Table 5).

### Speciation history of the *Pneumocystis* genus

These new complete genome data enabled us to examine the relationships between different *Pneumocystis* species and to estimate the timing of speciation events that led to the extant species. We inferred a strongly supported phylogeny of *Pneumocystis* species rooted with outgroups from distantly related fungal subphyla. Our phylogenomic analysis of 106 single-copy orthologs inferred from all assemblies including the fragmented Ck2 strongly supports monophyly of *Pneumocystis* species (100% Maximum likelihood bootstrap values; Fig. 2a), Bayesian posterior probabilities (>0.95; Supplementary Figure 1), and highly significant support from the Shimodaira-Hasegawa test (*18*) (*p* < 0.001; see Methods). An identical phylogeny was recovered using mitochondrial genome data from 33 specimens representing 7 *Pneumocystis* (Supplementary Figure 2). However, we identified unexpected placements of *P. wakefieldiae*, *P. oryctolagi* and *P. canis*. First, *P. wakefieldiae* appears as a sister species of *P. murina* instead of *P. carinii* (which also infects rats) (Fig. 2b). This observation is supported by the higher similarity in genome size (Supplementary Table 3), sequence divergence (Supplementary Table 4), genome structure (Fig. 1; Supplementary Table 5) and higher frequencies of supporting genes (0.64 in 1718 nuclear gene trees examined; Methods) between *P. wakefieldiae* and *P. murina* than between *P. wakefieldiae* and *P. carinii*. These relationships contradict the previous phylogenetic placement of *P. wakefieldiae* as an outgroup of the *P. carinii/P. murina* clade (*9*) or a sister species of *P. carinii* (*19*) based on analysis of mitochondrial large and small subunit rRNA genes (mtLSU and mtSSU). The new phylogeny also opposes the prevailing hypothesis for dynamics of host specificity and coevolution within the *Pneumocystis* genus, that is, *P. wakefieldiae* shares with *P. carinii* the same host species (*Rattus norvegicus*) and thus is expected to be more related to *P. carinii* than to *P. murina*.

Similarly, *P. oryctolagi* would be expected to be phylogenetically closer to rodent *Pneumocystis* than to primate *Pneumocystis*, consistent with the closer phylogenetic relationships of rabbits and rodents to each other than to primates (*20*) (Figs. 2a and 2b). In contrast, *P. oryctolagi* and *P. canis* are more closely related to primate *Pneumocystis (P. jirovecii* and *P. macacae*) than rodent *Pneumocystis* (Fig. 2a; Supplementary Figures 1 and 2; 100% of tree level support in 1,718 nuclear genes). The phylogenetic discrepancy between *P. oryctolagi* and its host (rabbit) suggests that host switching may have occurred in their distant history.

From whole-genome Bayesian phylogenetic estimates (see Methods), the common ancestor of all extant species of the genus emerged around 140 mya (confidence intervals: 180–101 mya; Fig. 2c; Supplementary Figure 1), with a separation of *Pneumocystis* and *Schizosaccharomyces* genera around 512 mya (CI: 822-203 mya) which is consistent with independent estimates of the origin of Taphrinomycota crown group at 530 mya (*21*). The *Pneumocystis* genus thereafter divided into two main clades, P1 consisting of *P. jirovecii, P. macacae, P. oryctolagi* and *P. canis*, and P2 consisting of species infecting rodents (*P. carinii, P. wakefieldiae* and *P. murina*) (Fig. 2b). Subsequent to the divergence of P1/P2, the clade P1 diversified through a series of speciation events leading either to new primate or carnivore species whereas P2 remained localized in rodents. We also found that the divergence time of *Pneumocystis* in the clade P1 predates that of their hosts, that is, the crown of rodent-rabbit-primate *Pneumocystis* is clearly more ancient than the corresponding superorder of mammals (Euarchontoglires) (Fig. 2c). The pattern in clade P2 is different as the divergence time estimates overlap with those of their hosts (Fig. 2c). On the basis of coalescent estimates, *P. jirovecii* began to split from *P. macacae* at ~62 mya (CI: 69-55 mya) which extended through the Cretaceous-Paleogene mass extinction event at 66 mya, but substantially predates the crown Catarrhini (human-macaque ancestor) of ~20 mya (CI: 24-17 mya; Fig. 2c; Supplementary Figure 1).

### High levels of population differentiation identified from *Pneumocystis* genomes support reproductive isolation

To understand the genomic divergence landscape of *Pneumocystis* populations, we performed genome-wide differentiation tests (F_ST_, relative population divergence) and nucleotide diversity (π) (Methods). These analyses used 32 genomic datasets, including 26 publicly available datasets in GenBank for *P. jirovecii, P. carinii* and *P. murina* and 6 datasets generated in this study for other four *Pneumocystis* species (Supplementary Note 1; Supplementary Table 2). Of note *Pneumocystis* organisms from macaque, rat and rabbit samples are from infected laboratory or domesticated animals (Supplementary Table 5), and thus do not represent true random representation of natural (e.g. wild) populations. We used a trained version LAST (*22*) to account for interspecies divergence during read mapping and ANGSD (*23*) to derive genotype likelihoods instead of genotypes. Since ANGSD’s F_ST_ requires outgroups, we analyzed interspecies divergence between *P. jirovecii, P. macacae* and *P. oryctolagi* populations using a sliding window approach of 5-kb and *P. carinii* as an outgroup species (*n* samples = 59). *P. murina* genomic divergence relative to *P. carinii* and *P. wakefieldiae* populations was estimated similarly using *P. jirovecii* as an outgroup species (*n* = 47). We found high levels of population differentiation among *Pneumocystis* specimens; 71.9% of the *P. jirovecii* genome had a Fixation index (F_ST_) > 0.8 compared to the closest species, *P. macacae*, while 90.2% of the genome had a F_ST_ > 0.8 compared to the extant species *P. oryctolagi* (Supplementary Figure 3). Similarly, 86.3% and 93.7% of the *P. murina* genome had a F_ST_ > 0.8 compared to *P. carinii* and *P. wakefieldiae*, respectively (Supplementary Note 1).

### Analyzing historical hybridization in *Pneumocystis* genus

Topology-based maximum likelihood analysis of 1,718 gene trees using PhyloNet (*33*) found no evidence of statistically significant signals for gene flow among species of clade P1 (*P. jirovecii, P. macacae, P. oryctolagi* and *P. canis*) (see Methods; Supplementary Figure 4), which indicates that these lineages were reproductively isolated throughout their evolutionary history, consistent with their isoenzyme diversity (*34*). In contrast, we found strong evidence of ancient hybridization in clade P2, possibly between *P. carinii* and *P. wakefieldiae* (Methods; Supplementary Figure 4), which may then have contributed to the formation of the *P. murina* lineage. We hypothesize that *P. murina* might have originated as a hybrid between ancestors of *P. carinii* and *P. wakefieldiae* in rats, and subsequently shifted to mice, possibly owing to the geographic proximity of ancestral rodent populations (for example in Southern Asia (*35*)), which is consistent with the fact that ecological fitting is a major determinant of host switch (*36*). The presumed physiological, cellular and/or immunological similarities among closely related rodent species might also have helped the same *Pneumocystis* species colonizing multiple closely-related rodent hosts (*10, 36*). The putative host shift might have been required because of negative selection against *P. murina* specifically in rats, possibly stemming from competition of low-fitness hybrids with parental species as is frequently observed in fungal pathogens (*37*). It is also interesting to note that earlier studies have suggested competition between *P. carinii* and *P. wakefieldiae* in rat colonies (*38*), and further, that *P. wakefieldiae* can no longer be detected in commercial vendors (Cushion et al. unpublished observation), while *P. carinii* can consistently be identified in laboratory rats. However, both *P. carinii* and *P. wakefieldiae* can be found, alone or together, in different species of *Rattus* in Southeast Asia (*10*).

### Gene families and metabolic pathways linked to host specificity

Gene annotations of *P. macacae* and *P. wakefieldiae* genomes was performed using RNA-Seq paired-end reads to guide *ab initio* gene predictions (Methods). *P. oryctolagi* and *P. canis* genomes were annotated using *ab initio* and homology-based predictions. The predicted protein-coding gene numbers are similar across *Pneumocystis* genomes and range from 3,211 in *P. wakefieldiae* to 3,476 in *P. canis* strain Ck1 (Supplementary Table 3). Nearly all predicted protein coding genes in *P. macacae* (96% of 3,471) and *P. wakefieldiae* (99% of 3,221) genomes have RNA-Seq support. Gene models present a complex architecture with 5.7 to 6.3 exons per gene. High representation of core eukaryotic genes in *P. macacae, P. oryctolagi, P. canis* and *P. wakefieldiae* provides evidence that these genomes are nearly complete and comparable in completeness to *P. jirovecii, P. murina* and *P. carinii* genomes: 86.2 to 93.4% of conserved genes are detectable in all annotated genome assemblies (Supplementary Table 3).

Examination of orthologous genes reveals that ~3,100 orthologous clusters had representative genes from all nine analyzed genome assemblies from seven *Pneumocystis* species (Supplementary Table 3). We found a small number of unique genes in each *Pneumocystis* species ranging from 25 in *P. wakefieldiae* to 204 in *P. oryctolagi* (Supplementary Table 3). Unique genes in most species encode for phylogenetically unrelated proteins with unknown function. A striking exception is observed in *P. macacae* in which nearly all unique proteins are part of a novel undescribed large protein family (*n* = 190). The members of this new family are enriched in arginine and glycine amino acid residues (denoted RG proteins) (Supplementary Figure 5a) and have no similarities with transposable elements. While RG motifs are often found in eukaryotic RNA-binding proteins (*24*), *P. macacae* RGs do not possess an RNA-binding domain (Pfam domains PF00076, PF08675, PF05670, PF00035), suggesting a different role. In addition, *P. macacae* RGs lack functional annotation except for two proteins that encode a Dolichol-phosphate mannosyltransferase domain (PF08285) and a leucine zipper domain (PF10259), respectively. Of the 190 RGs, 134 have RNA-Seq based gene expression support, including five among the top highly expressed genes (Supplementary Figure 5b). Nearly half of RGs are located at subtelomeric regions and often found in close proximity to *msg* genes (Supplementary Table 6). RG proteins can be grouped in three main clusters (based on OrthoFinder clustering; Methods), have a reticulate phylogeny (Supplementary Figure 5c) and a mosaic gene structure (Supplementary Figure 5d) which suggest frequent gene conversion events. These results suggest that RG proteins may play important roles in *P. macacae* specific biology. Further experiments are ongoing to elucidate the functions of these proteins in *P. macacae*.

To investigate the gene loss patterns in newly sequenced genomes, we compared *Pneumocystis* gene catalogs to those of related Taphrinomycotina fungi. We found that all sequenced *Pneumocystis* species have lost ~40% of gene families present in other Taphrinomycotina (Supplementary Figure 6), and that the metabolic pathways are also very similar among *Pneumocystis* species with a few minor (possibly stochastic) differences (Supplementary Note 2). This strongly suggests that *Pneumocystis* ancestry experienced massive gene losses that occurred before the genus diversification.

To investigate changes in gene content that might explain interspecies differences among the seven *Pneumocystis* species, we searched for expansions or contractions in functionally classified gene sets. We identified Pfam domains with significantly uneven distribution among genomes (Wilcoxon signed-rank test *p* < 0.05). Domains associated with Msg proteins are enriched in *P. jirovecii* and, to a lesser extent in *P. macacae* compared to other species (Fig. 3a). Domains associated with peptidases (M16) are enriched in *P. carinii, P. murina* and *P. wakefieldiae*. S8 peptidase family (kexin) is expanded in *P. carinii* as described previously (*13*) with 13 copies whereas all other species have one or three copies (Fig. 3a; Supplementary Figure 7). Although kexin is localized in other fungi to the Golgi apparatus, and in *Pneumocystis* is believed to be involved in the processing of Msg proteins, the expanded copies are predicted to be GPI-anchored proteins, appear to localize to the cell surface; their function is unknown (*25*). *P. carinii* and *P. wakefieldiae* have 13 and 3 copies whereas all other *Pneumocystis* species have only one (Supplementary Figure 7). We found that *P. carinii* kexin genes evolved under strong positive selection (*p* = 0.008) whereas *P. wakefieldiae* kexin genes did not (*p* = 0.159).

**Fig. 3.**
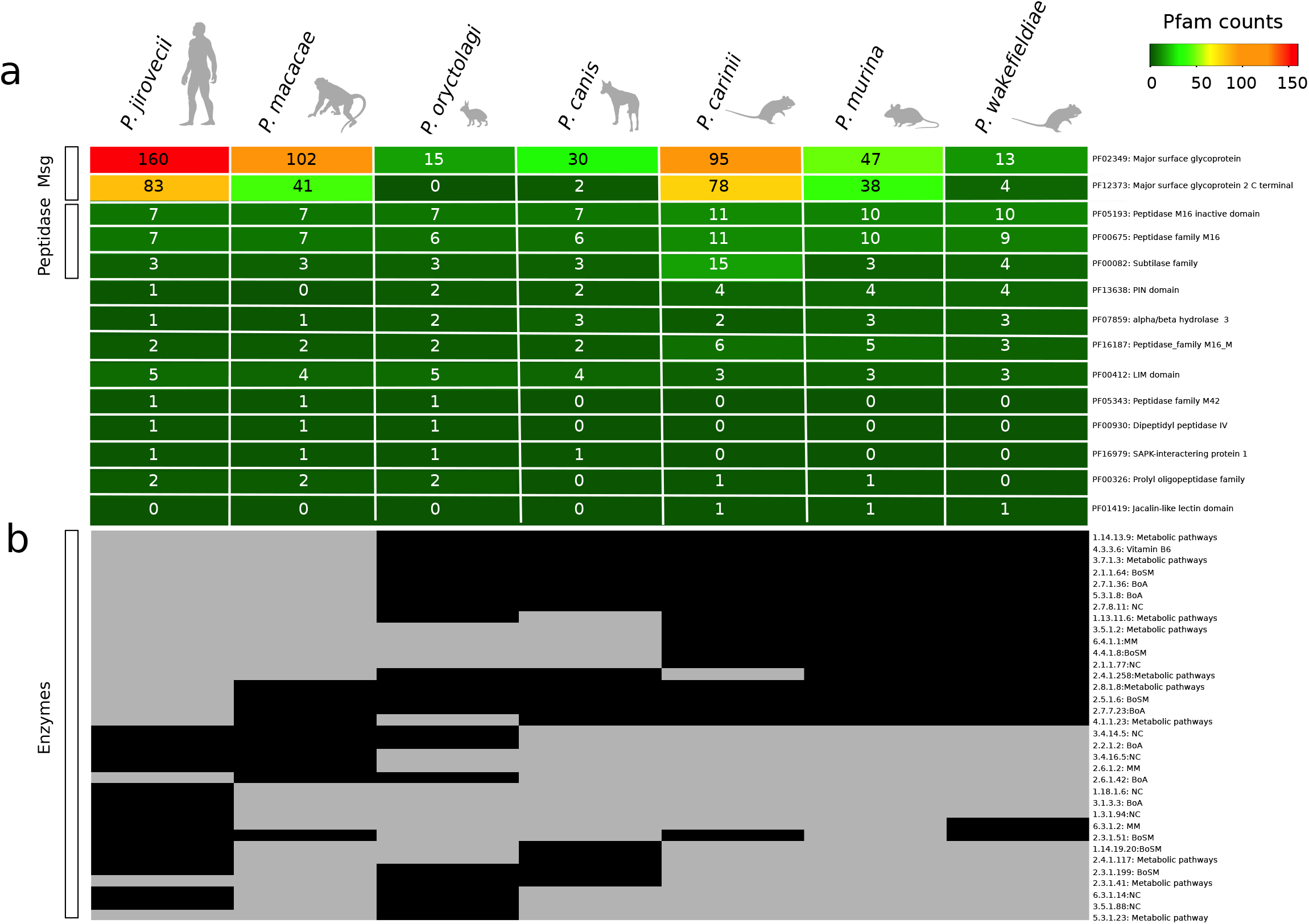
Distribution of protein families among *Pneumocystis* species. a, Heatmap of Pfam protein domains with significant differences (Wilcoxon signed-rank test, *p* < 0.05) are included if the domain appears at least once in the following comparisons: primate *Pneumocystis (P. jirovecii* and *P. macacae*) versus other *Pneumocystis*, clade P1 (*P. jirovecii, P. macacae, P. oryctolagi, P. canis* Ck1) versus clade P2 (*P. carinii, P. murina* and *P. wakefieldiae*), primate *Pneumocystis* versus clade P2. The number of proteins containing each domain is indicated within each cell for each species. The heat map is colored according to a score, as indicated by the key at the upper right corner. b, Heatmap of distribution of enzymes (represented by Enzyme Commission numbers), with their presence and absence indicated by black and grey colored cells, respectively.

Proteins with CFEM (common in fungal extracellular membrane) domains are important for the acquisition of vital compounds in fungal pathogens (*26*). *Pneumocystis* have an unusual high presence of CFEM domains compared to other Taphrinomycotina; each species possesses five proteins containing 2 to 6 domains per protein (Supplementary Figures 8a-c), with no significant differences among different species (*p*-value = 0.057; PF05730.10). Phylogenetic analysis of CFEM domains indicates that *Pneumocystis* species have experienced significantly higher intragenic duplications rates relative to other fungi (Supplementary Figure 8b). These results suggest that multiple CFEM domains were likely already present in the last common ancestor of *Pneumocystis* and were vertically transmitted.

To investigate changes in enzyme gene content that might account for inter-species differences among *Pneumocystis* species, we searched for enzymes that show clear differences among species, which are represented by Enzyme Commission numbers (ECs) (Fig. 3b). We found 34 ECs, which include 14 that are highly conserved in *P. jirovecii* but have a patchy distribution in other members of clade P1 (*P. macacae, P. canis* and *P. oryctolagi*) and are lost in clade P2 (*P. carinii, P. murina* and *P. wakefieldiae*). Most of these 14 ECs are assigned to the biosynthesis of antibiotics or secondary metabolites and vitamin B6 metabolism according to KEGG pathways. The latter pathway seems only functional in P2 clade (Supplementary Note 2).

### Intron evolution

We analyzed 1,211 one-to-one gene orthologs shared by all sequenced *Pneumocystis* and other Taphrinomycotina fungi (Supplementary Figure 9a). A total of 9,080 homologous sites within 1,211 alignments were identified (Supplementary Figure 9b). While intron densities are similar among *Pneumocystis* species (ranging from 4,842 in *P. macacae* to 5,289 in *P. murina*), they are markedly more elevated compared to related Taphrinomycotina, including *Neolecta irregularis (n* = 4,202 introns), *Schizosaccharomyces pombe* (*n* = 862) and *Taphrina deformans (n* = 639) (Supplementary Fig. 11b). Predictions of ancestral intron densities show that the *Pneumocystis* common ancestor had at least 5,341 introns, of which 37% were novel i.e. not found in other Taphrinomycotina (Supplementary Figure 9c). This is in contrast to other fungi; ~26% of *Neolecta* introns were independently acquired whereas *S. pombe* and *T. deformans* genomes have experienced significant intron losses, which is consistent with previous studies (*31, 32*). These results suggest the emergence of *Pneumocystis* genus was preceded by a significant amount of intron gain.

### Positive selection footprints in *P. jirovecii* genes

We tested the hypothesis that *P. jirovecii* has adapted specifically to humans after its separation with *P. macacae*, and that there will be footprints of directional selection in the genome that point to the molecular mechanisms of this adaptation. To infer *P. jirovecii*-specific adaptive changes, we compared the *P. jirovecii* one-to-one orthologs to those of *P. macacae* and *P. oryctolagi* using the branch-site likelihood ratio test (*39*). Positive selection was identified as an accelerated non-synonymous substitution rate. The test identified 244 genes (out of 2,466) with a signature of positive selection in the human pathogen *P. jirovecii* alone (Bonferroni corrected *p*-value < 0.05; Supplementary Table 7). Gene Ontology enrichment analysis of these genes with accelerate rates identified significant enrichment for the biological process “cellular response to stress” (adjusted using Benjamini-Hochberg *p*-value =1.9 x 10^-6^) and the molecular function “potassium channel regulator activity” (*p* = 2.8 x 10^-10^). Among the 244 genes, 197 are conserved in all *Pneumocystis* genomes available whereas 47 are absent in clade P2 only (*P. carinii, P. murina* and *P. wakefieldiae;* Fig. 2b). While the latter set of genes encode proteins of unknown function, analysis of Pfam domains shows a significant enrichment in the biological process “nucleoside phosphate biosynthetic” process (*p* = 9.9 x 10^-5^) and the molecular function “carbon-nitrogen lyase activity” (*p* = 2.8 x 10^-10^). Further investigations will be required to determine the precise functions of these genes.

### Subtelomeric gene families

Until recently, the only in-depth data on the subtelomeric gene families in *Pneumocystis* have come from the *P. jirovecii, P. carinii* and *P. murina* (*13, 40*). These genes, including *msg* and kexin, are believed to be important for antigenic variation, and are well represented in the assemblies of *P. macacae, P. oryctolagi, P. canis* and *P. wakefieldiae*. We provide a comprehensive analysis of their composition and evolution that complements our recent publication (*41*).

*P. macacae* subtelomeres encode numerous arrays of Msg and RG proteins (Supplementary Table 7). Phylogenetic analysis of adjacent genes revealed only a few instances of recent paralogs, which suggests that most of the duplications and subsequent positional gene arrangements are ancient. Three *P. macacae* subtelomeric regions have a nearly perfect synteny in *P. jirovecii* with the only difference being the absence of RG proteins in *P. jirovecii* (Supplementary Table 7). *P. oryctolagi* subtelomeres tend to be enriched in orphan genes that are not members of the Msg superfamily, and are of unknown function. *P. canis* subtelomeres are enriched in Msg-C family (see Msg section below). *P. wakefieldiae* subtelomeres are enriched in *msg* genes, though their types are distinct from those of *P. carinii* and *P. murina*.

### Evolution of *msg* genes

Up to 6% of the *Pneumocystis* genomes are comprised of copies of the *msg* superfamily, which are believed to be crucial mediators of pathogenesis through antigenic variation and interaction with the host cells. The superfamily is classified into five families A, B, C, D and E based on protein domain architecture, phylogeny and expression mode (*13, 40, 41*). The A family is the largest of the five, has been subdivided into three subfamilies (A1, A2 and A3) and is generally thought to contribute to antigenic variation. Their protein products contain cysteine-rich domain classified as N1 and M1 to M5.

To investigate the origin of *msg* genes, we used previously developed Hidden Markov Models (*13*) to search for corresponding gene models in the assemblies of *P. macacae, P. oryctolagi, P. canis* and *P. wakefieldiae* and combined these data with previously published *msg* sequences annotated in *P. jirovecii, P. carinii* and *P. murina* genomes (*13, 41*). Of note, in this study only a subset of *msg* genes were assembled for *P. oryctolagi, P. canis* and *P. wakefieldiae* due to difficulties in assembling highly similar short reads from Illumina sequencing exclusively while a potentially complete set of *msg* genes were assembled for *P. macacae* using Illumina and Nanopore reads (Supplementary Table 3). The number of full-length *msg* genes available ranges from 9 in *P. oryctolagi* to 161 in *P. jirovecii*. Sequence-based clustering and phylogenetic analyses of all *msg* genes (*n* = 482) revealed that: (i) there is no evidence of inter-species transfer among *Pneumocystis* species (Figs. 4b to d; Supplementary Figure 10), (ii) *msg* genes may have a polyphyletic origin, i.e. distinct families were present in most recent ancestors of *Pneumocystis* (Supplementary Figure 10a); (iii) *msg* genes experienced significant amount of recombination early in their history as estimated by phylogenetic network analysis (Supplementary Figures 10b and c).

**Fig. 4.**
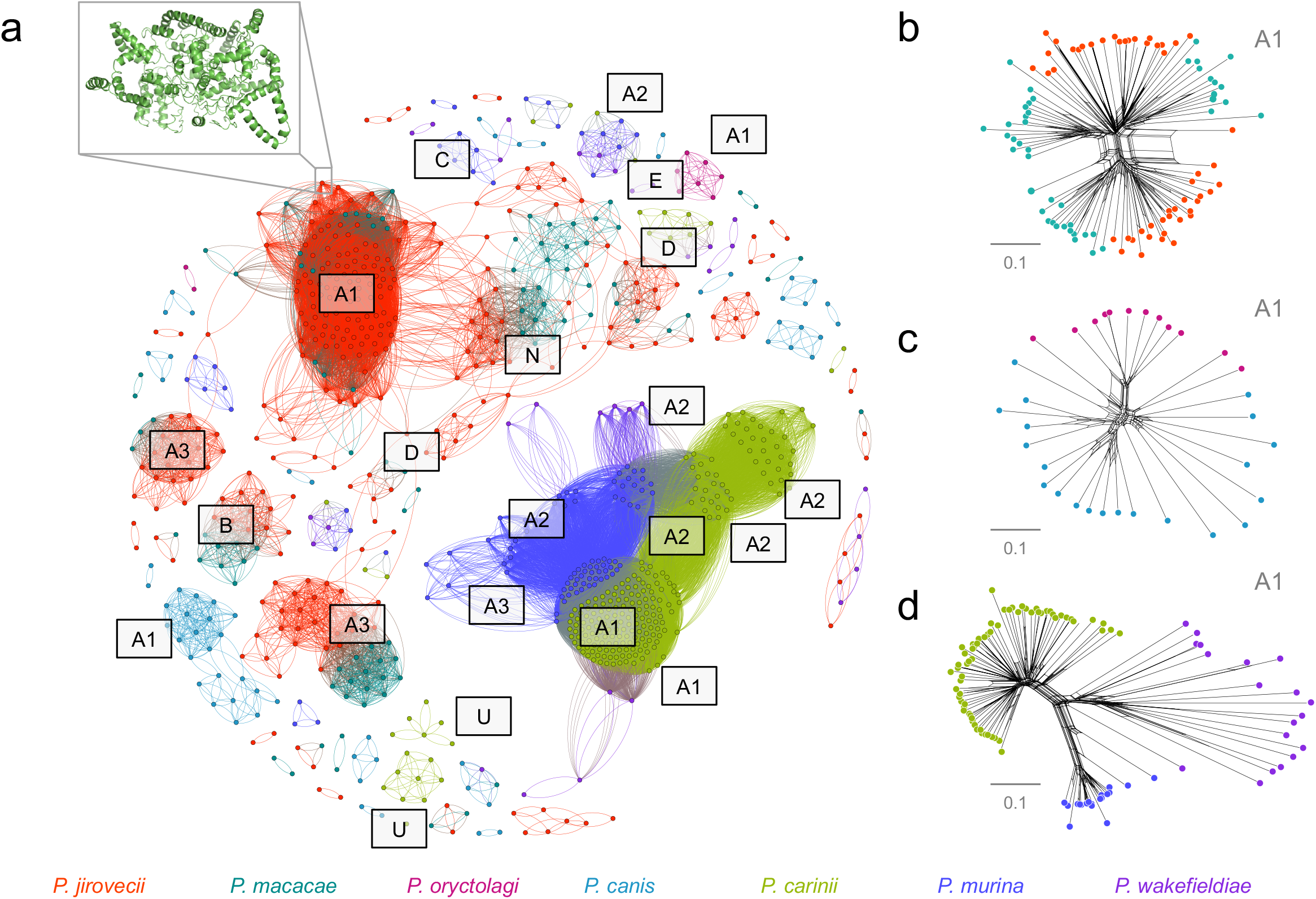
Clustering of *Pneumocystis* major surface glycoproteins (Msg). a, Graphical representation of similarity between 482 Msg proteins from 7 *Pneumocystis* species generated using the Fruchterman Reingold algorithm. A 3-D model of a representative Msg protein A1 family (NCBI accession number T551_00910) generated using DESTINI is presented in the upper left insert. Individual protein sequences are shown as dots and color-coded by species as shown at the bottom. The edge between two dots indicates a global pairwise identity equal or greater than 45%. The letters represent Msg families (A to E) and subfamilies (A1 to A3). N and U letters represent potentially novel Msg sequences (relative to our prior study (*41*)) and unclassified sequences, respectively. For sake of clarity only the major clusters were annotated. b, Phylogenetic network of a subset of Msg family A1 (*n* = 97) in primate *Pneumocystis* including *P. jirovecii* (red) and *P. macacae* (dark cyan) suggesting recombination events at the root of the network. Nodes with more than two parents represent reticulate events. Bar represent the number of amino acids substitution per site. c, Phylogenetic network of Msg family A1 (*n* = 33) in *P. oryctolagi* (red violet) and *P. canis* (light blue). d, Phylogenetic network of Msg family A1 (*n* = 113) in rodent *Pneumocystis* including *P. carinii* (green), *P. murina* (dark blue) and *P. wakefieldiae* (blue violet). The complete phylogenetic network is provided in Supplementary Data.

The evolution of *msg* genes between clades P1 and P2 is not uniform among *Pneumocystis* species and instead has clear differences between them. While some gene expansions are relatively recent (for example, *msg* families A, C and D) other expansions (*msg* families E and B) occurred before the emergence of *Pneumocystis* genus itself (Supplementary Figure 11). Subsets of *msg* genes show strong host specific sequence diversification (Fig. 4a), such as the current A family have emerged relatively recently 43 mya ago (CI: 58-28 mya) compared to the emergence of the genus at 140 mya ago (see Methods; Supplementary Figure 11). The A1 subfamily displays a substantial expansion in all species (Fig. 4a) and is subject to significant intra-species recombination (Figs. 4b to d), which suggest that *Pneumocystis* most recent ancestor may have develop a pre Msg-A family, which then evolved through duplication and recombination after the separation of species.

The A3 subfamily has expanded only in clade P1 (especially in *P. jirovecii*) whereas A2 has expanded only in clade P2 (*P. carinii, P. murina* and to a lesser extent in *P. wakefieldiae*) (Fig. 4a). Although all members of the A family might have a shared deep ancestry, we found no evidence suggesting that the A1, A2, A3 subfamilies are directly derived from one another (Supplementary Figure 10).

The *msg* B family underwent a net independent expansion in *P. macacae (n* = 10) and *P. jirovecii (n* = 12), while being reduced to only one copy in *P. oryctolagi* and *P. canis*, and being completely absent in *P. wakefieldiae*, *P. carinii* and *P. murina* (Fig. 4a). Using Bayesian estimates, we estimated the origin of B family to be older than the *Pneumocystis* genus itself (~211 versus 140 mya; Supplementary Figure 11). While a half of the B family members are located in subtelomeric regions in *P. jirovecii* and *P. macacae*, we found no evidence of recent inparalogs, which is consistent with their ancient origin. B family members lack predicted GPI anchor or transmembrane domain and have a shorter proline-rich motif compared to other *msg* families. Many of the *msg* B family members have a predicted secretory signal (13/25), with more in *P. jirovecii* than in *P. macacae* (7 vs. 3 copies). These data suggest that at least some members of the B family may be secreted effectors.

The *msg* D family is expanded only in *P. macacae* and *P. jirovecii*. The D family emerged at ~69 mya (CI: 109-40 mya) before the split of these two species (Supplementary Figure 11), thus suggesting a role in adaptation to primates similar to the A3 subfamily. In contrast, the E family, which is conserved in all species, is much more ancient at ~311 mya ago (CI: 541-158 mya), again preceding the emergence of the genus (Supplementary Figure 11).

*P. jirovecii* and *P. macacae* have a significantly larger number of *msg*-associated cysteine-rich domains than other *Pneumocystis* species (Fig. 5a) and also a much greater sequence diversity per domain than other *Pneumocystis* species (Fig. 5c). Domain sequences cluster independently, with each cluster containing sequences from all *Pneumocystis* species (Fig. 5b). Domains M1 and M3 are more closely related to each other than other domains, which suggests a relatively recent duplication. These results suggest that the all domains are likely to appear in the *Pneumocystis* ancestor and underwent a series of lineage specific expansions. The paucity of domains in *P. oryctolagi, P. canis* and *P. wakefieldiae* might reflect an interaction with host cells different than other species. Alternatively, these differences could represent incomplete sets of Msgs in the former species.

**Fig. 5.**
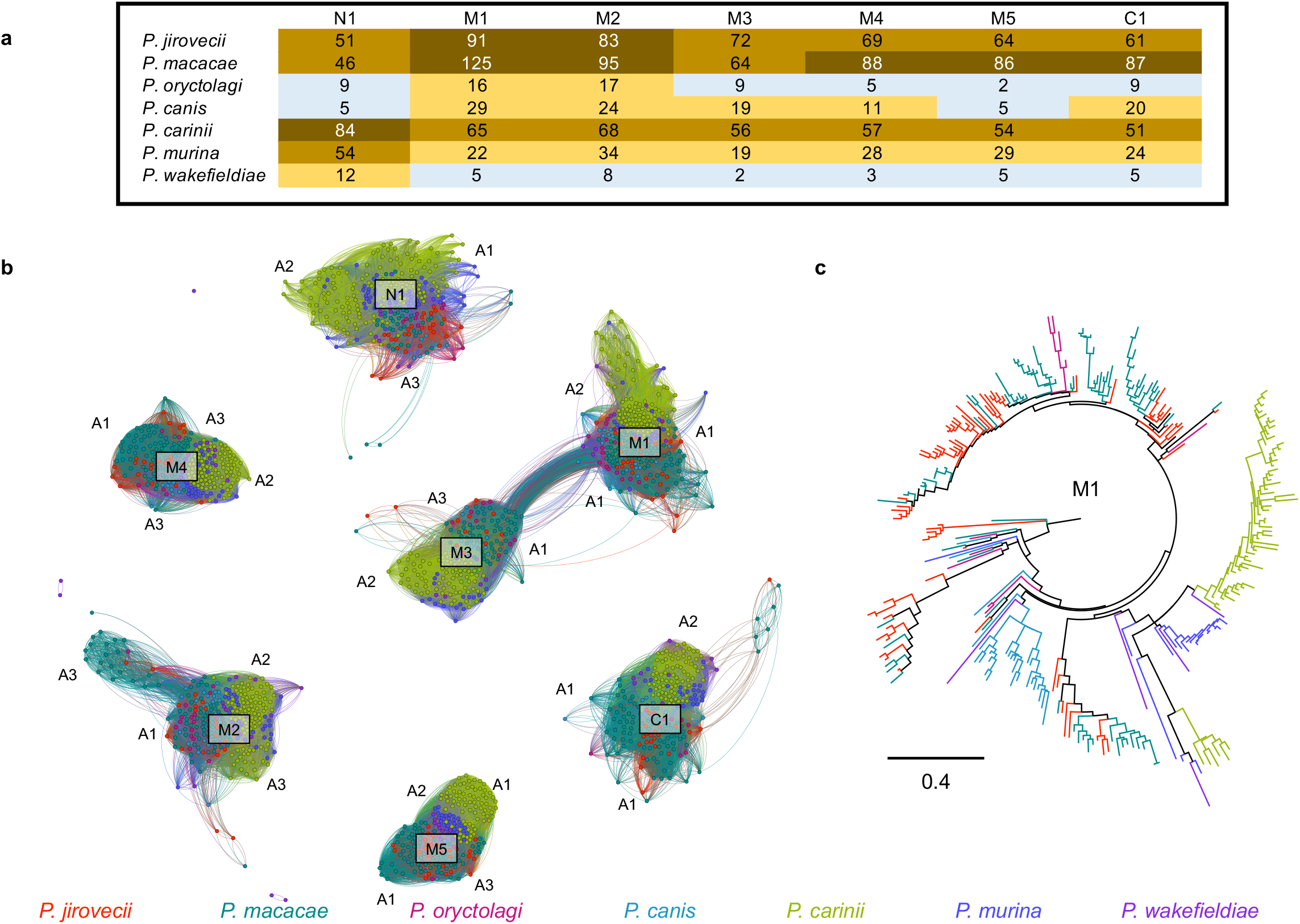
Evolution of Msg cysteine-rich protein domains in *Pneumocystis*. a, Heatmap showing the distribution of Msg domains in each *Pneumocystis* species. The color change from blue-orange-brown indicates an increase in the number of domains. b, Graphical representation of protein similarity between domains, which highlights that the domains were present in the most recent common ancester and were maintained other than perhaps domains M1 and M3. Domains are clustered by a minimum BLASTp cutoff of 70% protein identity. c, Maximum likelihood tree of the M1 domain. In both panels b and c, domains are color-coded by species as shown at the bottom.

### Conclusions

In the current study, we have produced high-quality genome assemblies and used them to investigate the speciation and host specific adaptation of multiple members of the *Pneumocystis* genus. We have established a robust phylogeny, presented genomic differences among species, identified a possible introgression among rodent-hosted *Pneumocystis* species, and discovered two phylogenetically distinct *Pneumocystis* lineages in dogs. Our analysis suggests that successful infection of humans by *P. jirovecii* has a deep evolutionary root accompanied by important genomic modifications. Surprisingly, analysis of core genomic regions of nuclear genomes did not identify clear differences that are suggestive of mechanisms for host-specific adaptation; instead it is the highly polymorphic multicopy gene families in subtelomeric regions that appear to account for this adaptation.

Based on our analysis, we propose the following series of events for the emergence and adaptation of *P. jirovecii* as a major human opportunistic pathogen (Fig. 6). First, there was a major shift of a *pre-Pneumocystis* lineage (possibly a soil- or plant-adapted organism) to mammals, which led to a significant genome reduction but with a significant proliferation of introns and expansions of cysteine-rich domain-containing proteins involved in immune escape and nutrient scavenging from hosts. *Pneumocystis* genomes encode multiple gene families that have experienced a rapid accumulation of mutations favoring fungal replication in mammals. Each *Pneumocystis* species has employed different strategies to adapt to their host including lineage-specific expansions of shared gene families such as *msg* A1, A3 and D in *P. jirovecii* or gain and expansion of RG proteins in *P. macacae*. In addition, some shared gene families also have acquired different properties (e.g., transmembrane domain and secreted signals) potentially contributing to host specificity. Our data point to the possibility that chromosomal rearrangements may play a role in the inhibition of gene flow between *P. jirovecii* and *P. macacae* leading to their speciation. The absence of a reliable culture method and the inability to genetically manipulate *Pneumocystis* prevents directly testing our model. Moreover, for the genes that we have now implicated in the process of host adaptation, only a few have been functionally characterized. Future studies on the role of these genes will be important to elucidate the molecular basis of in host specific adaptation by *Pneumocystis* pathogens.

**Fig. 6.**
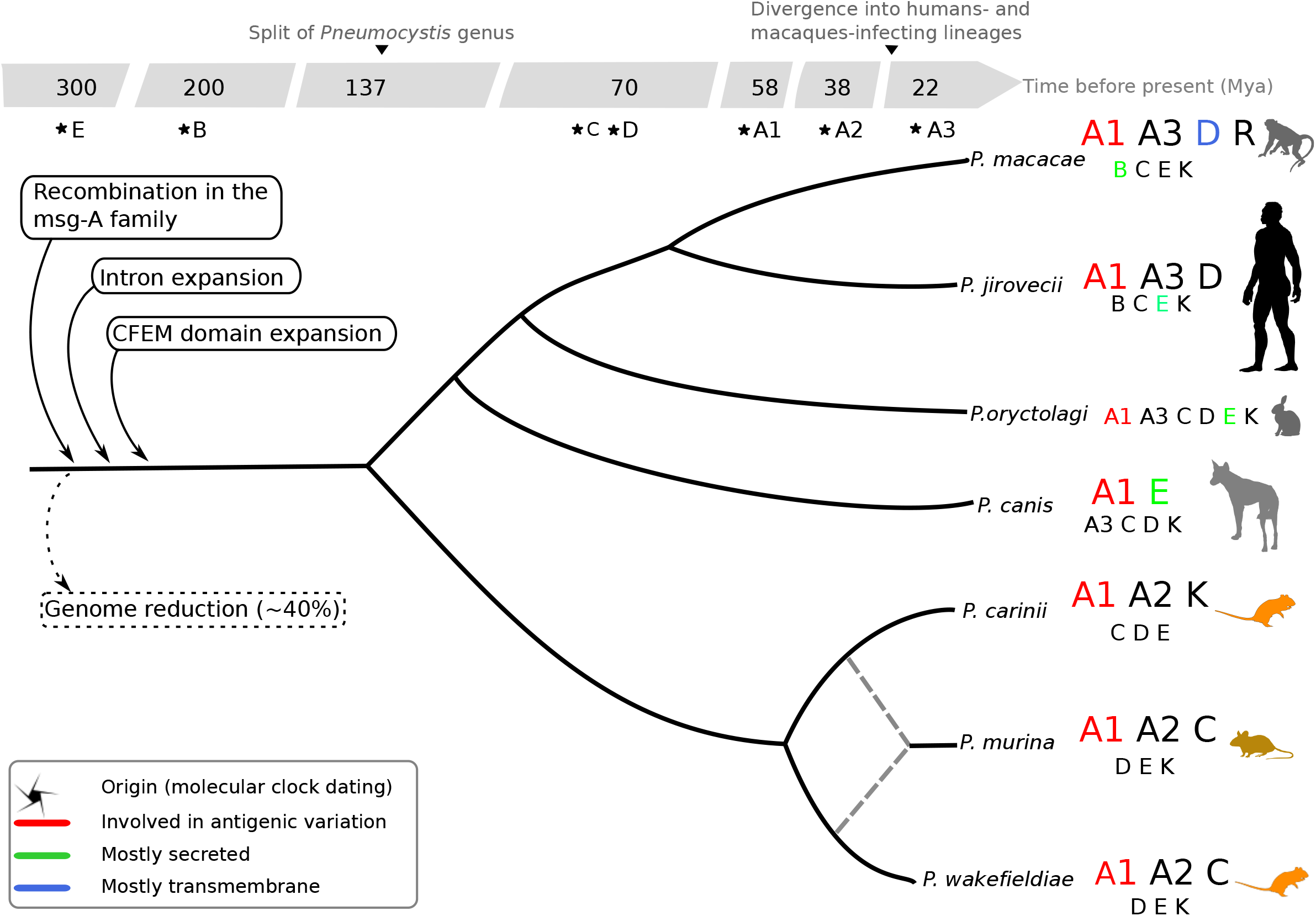
Overview of the genomic evolution of the *Pneumocystis* genus. Gene families are represented by letters: A to E for the five families of major surface glycoproteins (Msg) with the A family being further subdivided into three subfamilies A1, A2, and A3; K and R for kexins and arginine-glycine rich proteins, respectively. Larger fonts indicate expansions as inferred by maximum likelihood phylogenetic trees and networks. Dashed lines represent ancient hybridization between *P. carinii* and *P. wakefieldiae*. Detailed analysis also reveals distinct phylogenetic clusters within subfamilies. Introns and CFEM (common in fungal extracellular membrane) domains are enriched in *Pneumocystis* genes which indicate that these elements were likely present in the most recent common ancestor of *Pneumocystis* species. Animal icons were obtained from http://phylopic.org.

By untangling the co-evolution of *Pneumocystis* species with their mammalian hosts, we show that this evolution is more complex than portrayed by a strict co-evolutionary framework. The potential relaxation of strict host specificity in small mammals colonized by *Pneumocystis* could be explained as well by the fact that, in the coevolution theory, parasites infecting rodents (small-bodied with short lifespans, high reproduction rates, and high population densities) have lower host specificity than those adapted to long-lived large mammals with more stable population densities (*42*). Our analyses documented rare instances of pathogen speciation while sharing the same host (rats and dogs), which is equivalent to a speciation in sympatry (without geographical isolation). This work predicts novel and critical aspects of the genetic basis of host adaptation by *P. jirovecii*, the only fungal pathogen known to have adapted to living exclusively in human lungs. Future studies further expanding the sampled *Pneumocystis* genomes across the diversity of mammals, will be key to further understanding molecular basis of host specificity. The evolutionary processes that gave rise to the obligate biotrophic lifestyles of *Pneumocystis* within its host remain important future research questions (*43, 44*). The next important steps will also include the study of the influence of host biology on *Pneumocystis* adaptation, the genetic mechanisms underlying pathogen host shifts, and the genetic incompatibility between coexisting pathogens (e.g. *P. carinii* and *P. wakefieldiae* in rats).

## Material and Methods

### Experimental Design and *Pneumocystis* sample sources

Animal and human subject experimentation guidelines of the National Institutes of Health (NIH) were followed in the conduct of this study. Studies of human and mouse *Pneumocystis* infection were approved by NIH Institutional Review Board (IRB) protocols 99-I-0084 and CCM 19-05, respectively. The collection and processing of a single *P. jirovecii* human bronchoalveolar lavage sample from China (Pj55) was approved by the IRB of the First Affiliated Hospital of Chongqing Medical University, China (protocol no. 20172901). Written informed consent was obtained from the patient for the participation in this study. The authors confirmed that personal identity information of the patient data was unidentifiable from this report. The National Institute of Allergy and Infectious Diseases (NIAID) Division of Intramural Research Animal Care and Use Program, as part of the NIH Intramural Research Program, approved all experimental procedures pertaining to the macaques (protocol LVD 26). Nonhuman primate study protocols were approved by the Institutional Animal Care and Use Committee of the University of California, Davis (protocol no. 7092), the Tulane National Primate Research Center (TNPRC) and the Institutional Animal Care and Use Committee (IACUC) (protocol no. P0351R). Studies of rabbit *Pneumocystis* infection were reviewed and approved by the Institutional Animal Care and Use Committee of the University of Michigan (protocol no. RO00008218). For rabbit samples obtained France, the conditions for care of laboratory animals stipulated in European guidelines were followed (See: Council directives on the protection of animals for experimental and other scientific purposes, and J. Off. Communautés Européennes, 86/609/EEC, 18 December 1986, L358). Samples from *Pneumocystis* infected dog were collected as diagnostic samples and approved for only for research purpose. The owner’s consents for using samples and data were obtained on admission of the case and no further ethics permission was required because it was a routine diagnostic case and did not qualify as an animal experiment. Studies of rat *Pneumocystis* infection were approved by the Veteran Affairs animal protocol (VA ACORP #17-12-05-01). Clinical information and demographic data of the groups of individuals are presented in Supplementary Table 1.

Three *P. jirovecii* samples were obtained as bronchoalveolar lavage from patients at the NIH Clinical Center in Bethesda, MD, USA and Chongqing Medical University in Chongqing, China.

Six *P. macacae* samples were obtained as frozen lung tissues or formalin fixed paraffin embedded (FFPE) tissue sections prepared from SIV-infected rhesus macaques at the NIH Animal Center, Bethesda, Maryland (*n* = 2), the Tulane National Primate Research Center, Covington, Louisiana (*n* = 3), and the UC Davis California National Primate Research Center, Davis, California, USA (*n* = 1).

Four *P. oryctolagi* samples were obtained as frozen lung tissues from one rabbit with severe combined immunodeficiency at the University of Michigan, Ann Arbor, Michigan, USA, or as DNA from two corticosteroid treated rabbits and one rabbit with spontaneous *Pneumocystis* infection at the Institut Pasteur de Lille and the Institut National de la Recherche Agronomique de Tours Pathologie Aviaire et Parasitologie, Tours, France.

*P. canis* samples were obtained as DNA from one Cavalier King Charles Spaniel dog at the University of Helsinki, Finland and one Whippet mixed-breed at the University of Veterinary Medicine, Vienna, Austria.

*P. murina* organisms were obtained from heavily-infected CD40L-KO mice following a short-term *in vitro* culture. Genomic data obtained from *P. murina* isolates were combined with previously sequenced public data (Supplementary Table 2) and used for population genomics analysis (section “Speciation history of the *Pneumocystis* genus” and Supplementary Note 1).

One frozen cell pellet and 4 agarose gel blocks containing *P. wakefieldiae* and *P. carinii* were obtained from immunosuppressed rats (one gel block per rat) housed at the Cincinnati VA Medical Center, Veterinary Medicine Unit, Cincinnati, Ohio.

### Genome sequencing, assembly and annotation

Genomic DNA in agarose gel blocks was extracted using the Zymoclean Gel DNA Recovery Kit (Zymo Research). Genomic DNA in FFPE sections was extracted using the AllPrep DNA/RNA FFPE Kit (Qiagen). Genomic DNA in frozen lung tissues from two *P. macacae-infected* macaques and one *P. oryctolagi-infected* rabbit was extracted using the *Pneumocystis* DNA enrichment protocol as described elsewhere (*5, 13*). Genomic DNA in bronchoalveolar lavage samples from *P. jirovecii*-infected patients was extracted using the MasterPure Yeast DNA purification kit (Epicentre Biotechnologies, Madison, WI, USA). Total RNAs for *P. macacae, P. wakefieldiae* and *P. murina* were isolated using RNeasy Mini kit (Qiagen).

For DNA samples with small quantity, including three *P. oryctolagi* samples (RABF, RAB1 and RAB2B) and one *P. jirovecii* sample (RU817), we performed whole genome amplification prior to Illumina sequencing. Five microliters of each DNA sample were amplified in a 50-ul reaction using an Illustra GenomiPhi DNA V3 DNA amplification kit (GE Healthcare, United Kingdom).

Genomic DNA samples were quantified using Qubit dsDNA HS assay kit (Invitrogen) and NanoDrop (ThermoFisher). RNA samples integrity and quality were assessed using Bioanalyzer RNA 6000 picoassay (Agilent). The identities of *Pneumocystis* organisms were verified by PCR and Sanger sequencing of mtLSU before high throughput sequencing. For most of the DNA samples, at least one microgram of each DNA or RNA (depleted of ribosomal RNA using Illumina Ribo-Zero rRNA Removal Kit) sample was sequenced commercially using the Illumina HiSeq2500 platform with 150 or 250-base paired-end libraries (Novogene Inc, USA) or for one DNA sample of *P. jirovecii* from a Chinese patient using a single-read SE50 library using the MGISeq 2000 platform (MGI Tec, China). Raw reads statistics and NCBI SRA accession numbers are presented in Supplementary Material (Data access section) and Supplementary Table 2.

Adapters and low-quality reads were discarded using trimmomatic v0.36 (*45*) with the parameters “-phred33 LEADING:3 TRAILING:3 SLIDINGWINDOW:4:15 MINLEN:36”. Host DNA and other contaminating sequences were removed by mapping against host genomes using Bowtie2 v2.4.1 (*46*). Filtered Illumina reads were assembled *de novo* using Spades v3.11.1 (*47*). Details for host DNA sequences removal, *Pneumocystis* reads recovery and *de novo* assembly protocols are presented in Supplementary Material. Completeness of assemblies was estimated using BUSCO v9 (*48*), FGMP v1.0.1 (*49*) and CEGMA v2.5 (*50*).

Nanopore sequencing was performed on *P. macacae* DNA samples prepared from a single heavily infected macaque (P2C) with ~68% *Pneumocystis* DNA based on prior Illumina sequencing (Supplementary Table 2). High molecular weight genomic DNA fragments were isolated using the BluePippin (Sage Science) with the high-pass filtering protocol. A DNA library was prepared using the rapid Sequencing kit (SQK-RAD0004) from Oxford Nanopore Technologies (Oxford, UK) and loaded in the MinION sequencing device. Host reads were removed by mapping to the Rhesus macaque genome (NCBI accession number GCF_000772875.2_Mmul_8.0.1) using Minimap2 v2.10 (*51*). Unmapped reads were aligned to the draft version of *P. macacae* assembly built previously using Illumina data (Supplementary Methods) with ngmlr v0.2.7 (*52*). A total of 1,633,376 nanopore reads were obtained, of which ~5% were attributed to *Pneumocystis* (27-fold coverage), which is much less than the 68% based on Illumina data (Supplementary Table 2). This suggests that many *P. macacae* genomic DNA fragments were too short to pass the size selection filter. *Pneumocystis* nanopore reads were assembled using Canu v.1.8.0 (*53*), overlapped with the assembly using Racon v.1.3.3 (*54*) and polished with Pilon v1.22 (*55*) using the Illumina reads aligned with BWA MEM v0.7.17 (*56*).

Illumina RNA-Seq of the *P. macacae* sample P2C yielded 22 millions reads, of which ~92% were attributed to *Pneumocystis* (Supplementary Table 2). Filtered reads were mapped to the *P. macacae* assembly using hisat2 v2.2.0 (*57*), sorted with SAMtools v1.10 (*58*) and filtered with PICARD v2.1.1 (http://broadinstitute.github.io/picard). *De novo* transcriptome assembly of filtered reads was performed with Trinity (*59*). Quantification of transcript abundance was performed using Kallisto v0.46.1 (*60*). *P. wakefieldiae* (2A) and *P. murina* RNA-Seq data were processed similarly (Supplementary Table 2).

DNA transposons, retrotransposons and low complexity repeats were identified using RepeatMasker (*61*), RepBase (*62*) and TransposonPSI (http://transposonpsi.sourceforge.net). *Pneumocystis* telomere motif “TTAGGG” (*16*) was searched using “FindTelomere” (available at https://github.com/JanaSperschneider/FindTelomeres). The genomes of *P. carinii* strain Ccin (*14*) and strain SE6 (*12*) were scaffolded with Satsuma (*63*) using the *P. carinii* strain B80 as reference genome (*13*). *P. macacae, P. oryctolagi, P. canis* Ck1, *P. canis* A, *P. wakefieldiae*, and *P. carinii* (strains Ccin and SE6) genome assemblies were annotated using Funannotate v1.5.3 (DOI 10.5281/zenodo.1134477). The homology evidence consists of fungal proteins from UniProt (*64*) and BUSCO v9 fungal proteins (*48*). For *P. macacae* and *P. wakefieldiae*, RNA-Seq mapping files (BAM) and *de novo* transcriptome assemblies were used as hints for AUGUSTUS (*65*). *Ab initio* predictions were performed using GeneMark-ES (*66*). All evidences were merged using EvidenceModeler (*67*). *Taphrina* genomes (*T. deformans, T. wiesneri, T. flavoruba* and *T. populina* (*32, 68*)) and *P. canis* Ck2 were annotated using MAKER2 (*69*) because predicted gene models showed a better quality than those obtained from Funannotate. MAKER2 integrates *ab initio* prediction from SNAP (*70*), AUGUSTUS built-in *Pneumocystis* gene models (*71*) and GeneMark-ES as well as BLAST-based homology evidences from a custom fungal proteins database. GPI prediction was performed using PredGPI (*72*), big-PI (*73*) and KohGPI (*74*). Signal peptide leader sequences and transmembrane helices were predicted using Signal-P version 5 (*75*) and TMpred (*76*), respectively. Protein domains were inferred using Pfam database version 3.1 (*77*) with PfamScan (https://bio.tools/pfamscan_api). Domain enrichment analysis was performed using dcGOR version 1.0.6 (*78*). PRIAM (*79*) release JAN2018 was used to predict ECs using the following options: minimum probability > 0.5, profile coverage > 70%, check catalytic – TRUE and e-value < 10^-3^. *Pneumocystis* mitochondrial genome assembly and annotations are described in Supplementary Material. Three dimensional (3D) protein structure prediction of Msg proteins was performed using DESTINI (*80*) and visualized with PyMol (www.pymol.org).

### Comparative genomics

All genomes were pairwise aligned to the *Pneumocystis jirovecii* strain RU7 genome NCBI accession GCF_001477535.1 (*13*) using LAST version 921 (*22*) with the MAM4 seeding scheme (*81*). One-to-one pairwise alignments were created using maf-swap utility of LAST package and merged into a single multi-species whole genome alignment using LAST’s maf-join utility. Pairwise rearrangement distances in terms of minimum number of rearrangements were inferred using GRIMM (*82*) and Mauve (*83*). Breakpoints of genomic rearrangements were refined with Cassis (*84*) and annotated using BEDtools (*85*) ‘annotate’ command. Average pairwise genome-wide nucleotide divergences were computed with Minimap2 (*51*). Synteny visualization was carried out using Synima (*86*). Msg protein similarity networks were based on global pairwise identity obtained from pairwise alignments of full length proteins using Needle (*87*) or BLASTp (*88*) identity scores for individual protein domains. The networks presented in Figures 4 and 5 were generated using the Fruchterman Reingold algorithm as implemented in Gephi 0.9.2 (*89*).

To investigate the evolution of introns in *Pneumocystis* species, we identified unambiguous one-to-one orthologous clusters using reciprocal best Blast hit (e-value of 10^-10^ as cut off) in seven *Pneumocystis* species as well as in three other Taphrinomycotina fungi: *S. pombe, T. deformans* and *N. irregularis*. Intron position coordinates were extracted from annotated genomes using Replicer (*90*) and projected onto protein multiple alignments using custom scripts. Homologous splice sites in annotated protein sequence alignments were identified using MALIN (*91*). We required at least 11 unambiguous splicing sites and 5 minimal non-gapped positions. A potential splice was considered unambiguous if the site has at least 5 nongaps positions in the aligned sequences in both the left and right sides. MALIN uses a rates-across sites markov model with branch specific gain and loss rates to infer evolution of introns. Gain and loss rates were optimized through numerical optimizations. Fungi have a strong tendency to intron loss with few exceptions (e.g. *Cryptococcus*) whereas gain of intron is relatively rare. Thus, we penalized intron gain and set the variation rate to 4/3 for loss and gain levels. Intron evolutionary history was inferred using a posterior probabilistic estimation with 100 bootstrap support values.

### Phylogenomics

Orthologous gene families were inferred using OrthoFinder v.2.3.11 (*92*). In addition to *Pneumocystis* and *Taphrina* species, the predicted proteins for the following species were downloaded from NCBI: *Neolecta irregularis* (accession no. GCA_001929475.1), *S. pombe* (GCF_000002945.1), *S. cryophilus* (GCF_000004155.1), *S. octosporus* (GCF_000150505.1), *S. japonicus* (GCF_000149845.2), *Saitoella complicata* (GCF_001661265.1), *Neurospora crassa* (GCF_000182925.2), *Cryptococcus neoformans* (GCF_000149245.1), *Rhizopus oryzae* (GCA_000697725.1) and *Batrachochytrium dendrobatidis* (GCF_000203795.1). Single-copy genes were extracted from OrthoFinder output (*n* = 106) and concatenated into a protein alignment containing 458,948 distinct alignment patterns (i.e. unique columns in the alignment) with a gap proportion of 12.2%. Maximum likelihood tree analysis was performed using RAxML v 8.2.5 (*93*) with 1,000 bootstraps as support values. The LG model (*94*) was selected as the best amino acid model based on the likelihood PROTGAMMAAUTO in RAxML. 106 gene trees were estimated from each of the single copy genes. The Shimodaira-Hasegawa test (*18*) was performed on the tree topology for each of the gene trees and the concatenated alignment using IQ-Tree (*95*) with 1,000 RELL bootstrap replicates.

To infer the species phylogeny using mitochondrial genomes, protein coding genes were extracted, aligned using Clustal Omega (*96*), and concatenated. The resulting alignment was used to infer phylogeny using IQ-Tree v.1.6 (*97*) with TVM+F+I+G4 as the Best-fit substitution model and 1,000 ultrafast bootstraps and SH-aLRT test. A total of 33 mitogenomes from seven *Pneumocystis* species were used: *P. jirovecii [n* = 18 including 3 sequences from this study and four from previous studies (*5, 12, 17*)], *P. macacae (n* = 4), four *P. oryctolagi (n* = 4), *P. canis [n* = 4, (*17, 98*)], *P. carinii* [*n* = 2, (*17, 98*)], *P. murina* [*n* = 1, (*17*)] and *P. wakefieldiae* (*n* = 1).

Phylogenetic reconciliations of species tree and gene trees were performed using Notung (*99*). Ancestral reconstruction of gene family’s history was performed using Count (*100*). Phylogenetic network for Msg protein families was inferred using SplitTree (*101*). The detection of putative mosaic genes was performed using TOPALi v2.5 (*102*).

### Phylodating

Single-copy orthogroup nucleotide sequences were aligned using MACS v0.9b1 (*103*). Highly polymorphic *msg* sequences were excluded using BLASTn (*88*) with an e-value of 10^-5^ as cutoff against 479 published *msg* sequences (*13*). We inferred the divergence timing using two datasets: (1) 24 single-copy nuclear gene orthologs shared by all *Pneumocystis* and *S. pombe;* and (2) 568 nuclear genes found in all *Pneumocystis* species. BEAST inputs were prepared using BEAUTi v2.5.1 (*104*). Unlinked relaxed lognormal molecular clock models (*105, 106*) and calibrated birth-death tree priors (*107*) were used to estimate the divergence times and the credibility intervals. The substitution site model HKY was applied (*108*). Three secondary calibration priors were used: (i) *P. jirovecii/P. macacae* divergence with a median time of 65 mya as 95% highest posterior density (HPD) (*5*)), (ii) the emergence of the *Pneumocystis* genus at a minimum age of 100 mya (*4*), and (iii) the *Schizosaccharomyces* – *Pneumocystis* split at ~ 467 mya (*109*). For the dataset 2, the 568 gene alignments were concatenated in a super alignment with 568 partitions, with each partition defined by one gene. Gene partitions were collapsed using PartitionFinder v2.1.1 (*110*) with the “greedy” search to find optimal partitioning scheme. The alignment was split in three partitions in BEAST. Three independent runs for each dataset were performed separately for 60 million generations using random seeds. Run convergence was assessed with Tracer v1.7.1 (minimum effective sampling size of 200 with a burn-in of 10%). Trees were summarized using TreeAnnotator v.2.5.1 (http://beast.bio.ed.ac.uk/treeannotator) and visualized using FigTree v. 1.4.4 (http://tree.bio.ed.ac.uk/software/figtree) to obtain the means and 95% HPD. Host divergences were obtained from the most recent mammal tree of life (*6*), available at http://vertlife.org/data/mammals. The dating of fungal gene families was performed similarly.

### Population genomics

Sequence data sources and primary statistics are presented in Supplementary Table 2. Adapter sequences and low-quality headers of base sequences were removed using Trimmomatic (*45*). Interspecies reads alignment was performed using LAST (*22*) with the MAM4 seeding scheme (*81*). Alignments were processed by last-split utility to allow inter species re-arrangements, sorted using SAMtools v1.10 (*58*). Duplicates were removed using and PICARD v2.1.1. To compute the F_ST_ and nucleotide diversity (Watterson, pairwise, FuLi, fayH, L), we calculated the unfolded site frequency spectra for each population using the Analysis of Next Generation Sequencing Data (ANGSD) (*23*). Site frequency spectra was estimated per base site allele frequencies using ANGSD (*23, 111*). Hierarchical clustering was performed using ngsCovar (*112*). All data were formatted to fit a sliding windows of 1-10 kb using BEDTools (*113*). For each window, an average value of the statistics was calculated using custom scripts.

### Gene flow inference

To infer a phylogenetic network, we used 1,718 one-to-one orthologs from gene catalogs of seven *Pneumocystis* species using reciprocal best BLASTp hit with an e-value of 10^-10^ as cut off. Sequences from each orthologous group were aligned using Muscle (*114*). Alignments with evidence of intragenic recombination were filtered out using PhiPack (*115*) with a p-value of 0.05 as cut off. For each aligned group a maximum likelihood (ML) tree was inferred using RAxML-ng (*116*) with GTR+G model and 100 bootstrap replicates, and Bayesian tree was generated using BEAST2 (*104*). ML trees were filtered using the following criteria: 0.9 as the maximum proportion of missing data, 100 as the minimum number of parsimony-informative sites, 50 as the minimum bootstrap node-support value and 0.05 as the minimum p-value for rejecting the null hypothesis of no recombination within the alignment. BEAST trees with an effective sampling size < 200 were removed. Filtered trees were summarized using Treannotator (https://www.beast2.org/treeannotator/). Summary trees with an average posterior support inferior to 0.8 were discarded. Species network was inferred using PhyloNet option “InferNetwork_MPL” (*33*) with prior reticulation events ranging from 1 to 4. Phylogenetic networks were visualized using Dendroscope 3 (*117*).

The highest probability network inferred a hybridization between *P. carinii* and *P. wakefieldiae* leading to *P. murina* followed by a backcrossing between *P. murina* with *P. wakefieldiae* (log probability = −12759.4). Analysis of tree topology frequencies revealed that 64% of the trees were consistent with the topology of (*P. carinii, (P. murina, P. wakefieldiae)*), 28% with the topology of (*P. wakefieldiae, (P. carinii, P. murina)*) and 8% with the topology of (*P. murina*, (*P. carinii, P. wakefieldiae*)).

### Detection of positive selection

To search for genes that have been subjected to positive selection in *P. jirovecii* alone after the divergence from *P. macacae*, we used the branch-site test (*39*) as implemented in PAML (*118*) which detects sites that have undergone positive selection in a specific branch of the phylogenetic tree (foreground branch). A set of 2,466 orthologous groups between *P. jirovecii, P. macacae* and *P. oryctolagi* was used for the test. d_N_/d_S_ ratio estimates per branch per gene were obtained using Codeml (PAML v4.4c) with a free ratio model of evolution. This process identified 244 genes with a significant signal of positive selection only in *P. jirovecii* (d_N_/d_S_ > 1).

### Statistical analysis, custom scripts and figures

All custom bioinformatic analyses were conducted using Perl v5.26.0 (http://www.perl.org/) or Python v.3.6 (http://www.python.org) scripts. Pipelines were written using Snakemake v5.11.2 (*119*). Custom scripts and pipelines are available https://github.com/ocisse/pneumocystis_evolution. Statistical analyses were conducted in R version 3.3.2 (*120*). Phylogenetic trees with geological time scale were visualized using strap version 1.4 (*121*). Sequence motifs were visualized using WebLogo (*122*). Multi-panel figures were assembled in Inkscape (https://inkscape.org). Icon credit in Figure 6: Anthony Caravaggi under the license https://creativecommons.org/licenses/by-nc-sa/3.0/_(mouse), Sam Fraser-Smith (vectorized by T. Michael Keesey) (dog). https://creativecommons.org/licenses/by/3.0/. Anthony Caravaggi https://creativecommons.org/licenses/by-nc-sa/3.0/ (rabbit).

## Supplementary Materials

Supplementary Methods.

Data access.

Note S1. Population genomics analysis.

Note S2. Metabolic pathways.

Fig. S1. The maximum clade credibility tree of *Pneumocystis* species.

Fig. S2. Maximum likelihood phylogeny constructed using a concatenated dataset of 15 protein coding genes from 33 *Pneumocystis* mitochondrial genomes.

Fig. S3. Genome-wide scans for footprints of natural selection in *Pneumocystis*.

Fig. S4. Evidence of ancient gene flow in rodent *Pneumocystis* only.

Fig. S5. Evolution of arginine-glycine (RG) rich proteins in *P. macacae*.

Fig. S6. Heatmap showing gene family distribution in *Pneumocystis*.

Fig. S7. Expansion of kexin peptidase families in *Pneumocystis*.

Fig. S8. Evolutionary history of CFEM domains in *Pneumocystis*.

Fig. S9. Evolutionary history of introns in *Pneumocystis* and Taphrinomycotina fungi.

Fig. S10. RAxML phylogeny and phylogenic networks of Msg genes.

Fig. S11. Phylodating of major surface glycoproteins in *Pneumocystis*.

Table S1. Clinical information and demographic data of individual samples.

Table S2. Statistics and a posteriori classification of reads used in this study.

Table S3. Statistics of different *Pneumocystis* genome assemblies.

Table S4. Genome rearrangements among different *Pneumocystis* species.

Table S5. Pairwise nucleotide divergence (%) among *Pneumocystis* genomes.

Table S6. Subtelomeres in *P. macacae*

Table S7. *P. jirovecii* genome-wide signatures of selection.

## Author contributions

O.H.C, L.M and J.A.K conceived the project and designed all the experiments. L.M, O.H.C, C.W.L, J.B, J.X, J.S, R.B, B.P, K.V.R, R.K, A.S, M.C, V.H, J.C, L.P, M.T.C, G.K, Y.L, J.A.K performed the laboratory work to obtain samples for sequencing. O.H.C, L.M, J.P.D, P.P.K, J.L developed and implemented methods for sample processing, library preparation and sequencing. O.H.C, L.M, J.E.S, C.A.C, N.S.U analyzed the data. O.H.C, L.M and J.A.K drafted the manuscript, which was revised by all authors. J.E.S. and C.A.C. are CIFAR Fellows in the program Fungal Kingdom: Threats and Opportunities.

## Data and materials availability

All data needed to evaluate the conclusions in the paper are present in the paper and/or the Supplementary Materials.

The content of this publication does not necessarily reflect the views or policies of the Department of Health and Human Services, nor does mention of trade names, commercial products, or organizations imply endorsement by the U.S. Government.

