## Supplementary Materials for "Genomic insights into the host specific adaptation of the *Pneumocystis* genus and emergence of the human pathogen *Pneumocystis jirovecii*"

**This PDF file includes:**

Supplementary Materials and Methods

Data Access

Supplementary Notes 1 and 2

Supplementary Figures 1 to 11

Supplementary Tables 1 to 5

Supplementary References

**Other Supplementary Materials for this manuscript include the following:**

Supplementary Tables 6 and 7 (separate Excel spreadsheets)*.*

**Supplementary Materials and Methods**:

*P. macacae* genome sequencing: Host specific Illumina reads were removed by mapping against Rhesus macaque genome (NCBI accession number GCF_000772875.2_Mmul_8.0.1) using Bowtie2 (*1*) with the --sensitive-local option. Unmapped reads were merged and analyzed using GenomeScope (*2*) to estimate whether coverage was sufficient to successfully recover the genome. Genome assembly was performed using Spades v3.11.1 (*3*) and resulting scaffolds were aligned to the macaque genome using BLAT (*4*) with default parameters to remove residual contaminant contigs. Bacterial contaminants were removed by comparing the remaining scaffolds to NCBI nr database using BLASTx (*5*). Remaining scaffolds were compared to a database containing the published genomes of *P. jirovecii*, *P. carnii* and *P. murina* (GCA_001477535.1, GCA_001477545.1 and GCF_000349005.1) using BLAT with a minimum score of 200 nucleotides and a minimal identity of 70%. Scaffolds without significant similarity with *Pneumocystis* genomes as determined by BLAT were compared to a custom database of fungal proteins using TBLASTn (*5*) with an e-value of 0.01 as threshold. Mitochondrial contigs were identified by BLASTn against published *Pneumocystis* mitogenomes and removed.

*P. wakefieldiae* genome sequencing: *P. wakefieldiae* specimens were obtained from five rats (*Rattus norvegicus*) including three brown Norway rats and two Long Evans breed rats. Rats were acquired from different commercial vendors and housed at the University of Cincinnati from September 1995 to September 2002. *Pneumocystis* pneumonia was induced by corticosteroids-induced immunosuppression. All rats were co-infected by both *P. wakefieldiae* (NCBI taxonomy ID: 38082) and *P. carinii* (ID:4754) (Supplementary Table 1). We sequenced one genomic DNA preparation from each of the four rats and one sample of total RNA preparation from a single rat (Supplementary Table 2).

Raw Illumina HiSeq reads were filtered using trimmomatic (*6*) and aligned to the *Rattus norvegicus* genome (NCBI accession no. GCF_000001895.5) using Bowtie2 v.2.2.5 with the following parameters –very-fast –no-discordant –no-mixed. Unmapped reads were mapped against the *P. carinii* genome (NCBI accession # GCA_001477545.1) using Bowtie2 version 2.2.5 with the following parameters: -no-discordant -D 5 -R 1 -N 0 -L 32 -i S 0,2.50 –end-to-end. Reads that failed to map to Rat or *P. carinii* genomes were considered for initial genome assembly using Spades (*3*). We initially attempted to generate an assembly per isolate. However, this strategy yielded highly fragmented assemblies indicating insufficient coverage per sample. Of note, the removal of presumed polymorphic major surface glycoproteins (*msg*) gene reads before assembly did not substantially improved the assembly continuity or completeness (data not shown). Therefore, *msg*-related reads were not specifically removed before assembly in the subsequent steps.

Subsequently, filtered reads (i.e. after removing of high confidence rat and *P. carinii* reads from all samples) from all four rats were merged and assembled with Spades. This assembly process yielded 139,106 contigs totalizing 77 Mb, which were aligned to rat genome (GCF_000001895.5) using BLAT version 3.5 (*4*) with default parameters. Small sized scaffolds (< 500 bp) were filtered out because they appeared to be enriched in rodent specific repeats. This reduced the dataset to 10,630 contigs corresponding to 23 Mb.

To test whether our assembly truly captured *P. wakefieldiae* but not *P. carinii*, we predicted 7,918 gene models from the 10,630 scaffolds using AUGUSTUS version 3.2.1 (*7*) with built-in *Pneumocystis* gene models (*8*). We then performed an all-vs-all search using reciprocal BLASTp hit search with an e-value of 10^-10^ as threshold against a custom database containing the complete proteomes of *P. carinii* strain B80 (*n* = 3,646; NCBI accession no. GCF_001477545.1), *P. carinii* strain Ccin (*n* = 3,305; genome assembly from (*9*) but annotated in this study), *P. carinii* strain SE6 (*n* = 3,506; genome assembly from sample BALE6 (*10*) but annotated in this study) and *P. murina* (*n* = 3,838 proteins; GCF_000349005.1). Nucleotide sequence comparisons of seven homologous genes showed 4-7% divergence between *P. wakefieldiae* and *P. carinii* whereas only 0 – 0.8% divergence was observed within *P. carinii* species (*11*). Whole genome alignment-based divergence estimates between strains *P. carinii* can reach 5% (Supplementary Table 4), although this estimate may be inflated by higher error rates in strains Ccin and SE6. We BLASTed all the predicted genes against the *P. carinii* and *P. murina* reference gene sets (e-value of 10^-20^ as cut off). Candidate homologs were extracted and global pairwise identities were computed using Needle from EMBOSS package (*12*). Genes with at least 80% similarity with *P. carinii* homologs were extracted and analyzed. No gene was found with an identity > 80% with *P. murina*. For each gene, an orthologous group including sequences from *P. jirovecii*, *P. carinii* (strains B80, Ccin, SE6), *P. murina* and *S. pombe* when possible was constructed using reciprocal best BLASTp hit with an e-value of 10^-10^ as cut off. Multiple sequence alignments were generated using MAFFT (*13*) and phylogenetic inferences made with RAxML (*14*)*.* Based on phylogenetic placements and pairwise identities, 2,710 gene models located in 10,380 scaffolds were considered as *P. carinii* and removed. To remove residual non-coding *P. carinii* genomic segments, the remaining scaffolds (*n* = 250, size= 13,423,939 bp, GC=28.9%) were aligned to *P. carinii* strain B80 genome (GCA_001477545.1) with MegaBLAST (*15*) using the following parameters “-W 1000 -v 1 -b 1 -e 1e-150”. A total of 20 scaffolds were flagged by MegaBLAST and discarded after manual inspection, which left 230 scaffolds (7,200,860 bp, GC% = 30.1). We ran AUGUSTUS on these 230 scaffolds and clustered to predictions with the reference proteomes of *P. jirovecii*, *P. carinii* and *P. murina*. We identified 674 one-to-one orthologs, which were concatenated and used to build a phylogenetic tree using RAxML. This tree unambiguously placed *P. wakefieldiae* as distinct species from *P. carinii* and *P. murina*. After this validation step, we performed a gap closure and reduction using Redundans (*16*), which reduce the number of scaffolds to 134 totalizing 7.1 Mb (GC%=30).

We annotated these 134 scaffolds with AUGUSTUS and compared gene models to four published gene sequences of *P. wakefieldiae* sequences from Uniprot: Q01706 for guanine nucleotide binding protein alpha subunit, O00053 for PrBiP, O94108 for heat shock protein 70B/SSB1 (Fragment) and Q12652 for TATA binding protein. The PrBiP and TATA binding protein genes were fully recovered using reciprocal Best BLASTp hit with an e-value of 10^-10^ as cut off. The pairwise amino acid identities of our gene models with published PrBiP and TATA were 98.8% and 98.7%, respectively, whereas the identities with *P. carinii*, *P. murina* and *P. jirovecii* ranged from 82.8% to 96.5%. As part of the verification process, we performed a synteny analysis, comparing our assembly to the genome sequence of *P. carinii* and *P. murina* using SatsumaSynteny2 (*17*). We found that the gene order in our assembly was clearly different than orthologs in *P. carinii* and *P. murina*, which suggests a different species. To further verify the identity of our genome assembly, we identified the rRNA operon using BLAT and compated to the published *P. wakefieldiae* rRNA operon (GenBank under the accession no. L27658). We also identified a genomic fragment containing the mating-type locus cloned by PCR and Sanger sequenced in this study. While the rRNA operon was only partially recovered in our assembly (416 nt), the recovered part of the rRNA showed 99.8% nucleotide identity with the published sequence. The rRNA operon contains internal repeats which likely causes the assembly breakpoints. The fragment containing the mating-type locus (~15 kb) was fully recovered and exhibited 100% nucleotide identity. All internal gaps were closed by PCR and Sanger sequencing. The final genome assembly has 17 scaffolds with a total size of 7.3 Mb, which appears to be consistent with the results of karyotypic studies (*11*). This assembly clearly represents *P. wakefieldiae* based on sequence alignment and phylogenetic analysis of the four published genes of *P. wakefieldiae* described above (data not shown).

*P. canis* genome sequencing: We sequenced two *P. canis* DNA samples from two dogs from Austria and Finland (denoted as A and Ck, respectively; Supplementary Table 1). Since previous studies of the mitochondrial large subunit rRNA gene (mtLSU) have demonstrated the presence of two types of *P. canis* populations with a significant variability in the dog Ck (*18*), we analyzed the sequencing data from samples A and Ck separately. Two genome assemblies were recovered from the sample Ck (denoted as Ck1 and Ck2) and one assembly from the sample A. Due to the lack of reliable karyotype data, it remains unclear how many chromosomes there are for *P. canis*.

Prior to the genome assembly, the filtered reads were analyzed using GenomeScope (*2*) to determine if sufficient reads are available for full genome recovery. Reads were filtered using trimmomatic (*6*). Host reads were removed by mapping to dog reference genome (NCBI accession no. GCF_000002285.3) with Bowtie2 (*1*). In the dog Ck, preliminary assembly of these reads revealed the presence of exogenous DNA from the plant *Arabidopsis thaliana* and the fungus *Puccina graminis*. These reads were considered contaminants and removed by mapping reads against their respective genomes (GCF_000001735.3_TAIR10 and GCF_000149925.1). No contaminant was observed in reads from the dog A other than DNA from the host. After filtering, reads from each sample were assembled separately using Spades (*3*). We BLASTed the resulting scaffolds against UniProt bacterial database (available at [ftp.uniprot.org/databases/uniprot](ftp://ftp.uniprot.org/databases/uniprot)) for bacterial contamination using BLASTx (*5*) with an e-value of 10^-15^. Residual dog contaminants (mostly microsatellites) were detected by screening against a database of 96 *Canis familiaris* clone CSac3 satellite sequences downloaded from NCBI. Redundant and heterologous scaffolds were identified using Redundans (*16*). Haplotype analyses were performed using custom Perl scripts, which allow the identification of two populations.

*P. oryctolagi* genome sequencing: organisms were isolated from three wild rabbits collected in France and a single Interleukin-2 receptor-ɣ knockout rabbit with a severe combined immunodeficiency, from Michigan (*19*) (Supplementary Table 1). We sequenced a single genomic DNA specimen for each rabbit. The proportion of *Pneumocystis* DNA relative to host DNA was relatively low (< 5%; Supplementary Table 2). Filtered reads from all four samples were combined to generate a consensus genome assembly.

Illumina reads from the host were removed by mapping against rabbit genome (NCBI accession no. GCF_000003625.3_OryCun2.0) using Bowtie2 with the --sensitive-local option. Separate assembly of unmapped reads from each *P. oryctolagi* isolate resulted in a highly fragmented assemblies (for example, a total of 3.7 Mb in 2,645 contigs for the isolate RAB_F). Therefore, we pooled unmapped reads from all isolates and assembled them with Spades v3.11.1 (*3*). Initial assembly provided 1,544,250 contigs totalizing 877 Mb. A second round of filtering was performed comparing all contigs to rabbit genome using BLAT. After removal of host contigs, the remaining scaffolds were compared to a database containing the published genomes of *P. jirovecii*, *P. carinii* and *P. murina* (NCBI accession nos. GCA_001477535.1, GCA_001477545.1, GCF_000349005.1) using BLAT with a minimum score of 200 and a minimal identity of 70%. Scaffolds with no significant similarity with *Pneumocystis* genomes were compared to a custom database of fungal proteins using TBLASTn with an e-value of 0.01 as threshold. A total of 321 contigs totalizing 7.2 Mb passed that filtering step. To test if heterologous scaffolds were present in the assembly, we analyzed the 321 contigs using Redundans. Only 18 heterologous contigs were detected, of which 17 are smaller than 500 bp in size. The reduced assembly produced by Redundans pipeline was not used in subsequent analyses. We used “chromoAssemble” module of Satsuma2 (*17*) and *P. jirovecii* strain RU7 genome as reference to guide the synteny-based scaffolding. Genome annotation was performed using Funannotate (<https://zenodo.org/record/2604804>). The annotated heat shock protein 70 gene displayed 96% of nucleotide identity with the published sequence (DQ435616 (*20*)), which indicates some strain variation within *P. oryctolagi* populations.

*Pneumocystis* mitochondrial genomes assembly and annotation: Mitogenome reads were retrieved from Illumina HiSeq reads by mapping against published mitogenomes of *P. jirovecii, P. carinii* and *P. murina.* Mitogenome annotation was performed using the MFannot tool with genetic code 3 for yeast mitogenome (<http://megasun.bch.umontreal.ca/cgi-bin/mfannot/mfannotInterface.pl>). Transfer RNAs were further evaluated using tRNAscan-SE (*21*). Resulting open reading frames were reviewed and revised by comparing to an in-house database of published mitogenome genes in *P. jirovecii, P. carinii* and *P. murina* (114) using SeqMan Pro (version 14.1.0.118, DNASTAR, Madison, WI) under default conditions with a minimum match percentage of 70%. Retrieved reads were *de novo* assembled using the Sequencher software (version 5.4.6; Gene Codes Co., MI). The resulting contigs were realigned with Illumina raw reads using SeqMan Pro under default conditions with a minimum match percentage of 97%. Contigs not supported by raw reads were removed. The remaining contigs were assembled using Sequencher. For *P. wakefieldiae*, retrieved mitogenome reads were first aligned to the *P. carinii* mitogenome (GenBank accession number JX499145) with a minimum match percentage of 97%. Unaligned reads were *de novo* assembled using Sequencher. The final assembly of *P. wakefieldiae* mitogenome was amplified as 6 overlapping fragments from genomic DNA by PCR using primers specific for *P. wakefieldiae*. Selected regions from each PCR product were sequenced directly by Sanger Sequencing.

**Data Access:**

Genome assemblies and raw sequences data are available at: *P. macacae* (https://dataview.ncbi.nlm.nih.gov/object/PRJNA632025?reviewer=aj13lf1srqsu4l4t3jsd489m0m); *P. oryctolagi* (https://dataview.ncbi.nlm.nih.gov/object/PRJNA632560?reviewer=m2t967mnld23ipe986jed414n8); *P. canis* Ck1 (<https://dataview.ncbi.nlm.nih.gov/object/PRJNA632556?reviewer=jlk8amkh7t9lpklhn6kuk6k8i>); *P. canis* Ck2 (<https://dataview.ncbi.nlm.nih.gov/object/PRJNA632878?reviewer=728teg4f6gmev9u0o7u7dro09n>);

*P. canis* A (https://dataview.ncbi.nlm.nih.gov/object/PRJNA636786?reviewer=lctihg2er4g3r7v7sooh70lqud)*;*

*P. wakefieldiae* (https://dataview.ncbi.nlm.nih.gov/object/PRJNA632570?reviewer=gl29m28tg2877tl40nr8309efb).

*Pneumocystis jirovecii* strain 55 (PRJNA647920), *P. jirovecii* strain 54c (PRJNA648092), *P. jirovecii* strain 46 (PRJNA648096), *P. macacae* strain CJ36 (PRJNA648103), *P. macacae* strain ER17 (PRJNA648108), *P.* *macacae* strain UC86 (PRJNA648112), *P.* *macacae* strain GL92 (PRJNA648115). Individual SRA files are presented in Supplementary Table 2.

**Supplementary Text:**

**Note 1:** Population genomics analysis

Analysis of *P. jirovecii*, *P. macacae*, *P. oryctolagi*:

Coevolution of pathogenic species with their hosts leaves genomic footprints which can, if the divergence times are recent enough, be identified and characterized. These regions which are often referred as “genomic islands of differentiation” can reveal insights into the interaction of the pathogen with its host. We used genome scans to identify regions with significant divergence relative to average genomic divergence. We found that the *Pneumocystis* species genomes display a high level of genetic differentiation and are mostly evolving under weak purifying selection or neutral evolution. Genomic islands could not be identified because they have been eroded over the large evolutionary timescale that separates different *Pneumocystis* species. Specifically, to understand the genomic divergence landscape of *Pneumocystis* populations, we performed genome-wide differentiation tests (F_ST_, relative population divergence) and nucleotide diversity (π). We used a trained version LAST (*22*) to account for interspecies divergence during read mapping and ANGSD (*23*) to derived genotype likelihoods instead of genotypes. Since ANGSD’s F_ST_ requires outgroups, we analyzed interspecies divergence between *P. jirovecii*, *P. macacae* and *P. oryctolagi* populations using a sliding window approach of 5-kb and *P. carinii* as an outgroup species (*n* samples = 59). *P. murina* genomic divergence relative to *P. carinii* and *P. wakefieldiae* populations was estimated similarly using *P. jirovecii* as an outgroup species (*n* = 47). To analyze the genomic landscape of *P. jirovecii*, we aligned Illumina reads from *P. macacae*, *P. oryctolagi* and *P. carinii* to the *P. jirovecii* reference genome (strain RU7). Figure S3a illustrates the distribution of each isolate according to cluster distance. Hierarchical clustering unambiguously separates isolates from different species and indicates that the majority of genetic variation within this data set consists of inter-species fixed differences. F_ST_ scans reveal a differentiation between *P. jirovecii* and *P. macacae* populations and an even greater divergence with *P. oryctolagi*. In both comparisons, the genome-wide distributions of F_ST_ are unimodal with highest peaks at 0.98 and 0.93 (Supplementary Figure 3b). In fact, 71.9% of the total 16,825 windows of 5 kb are highly differentiated (F_ST_ > 0.8) in *P. jirovecii*-*P. macacae* comparison, and this number reaches 90.2% in *P. jirovecii-P. oryctolagi* comparison. Regions of F_ST_ < 0.5 are negligible 0.5% and 0.0%, respectively. Inspection of genomic regions with low F_ST_ reveals that they are significantly enriched in regions encoding variable major surface glycoproteins (Msg) (data not shown), which suggest that the low F_ST_ values are caused by local increases in genetic diversity.

Analysis for the trio *P. murina*, *P. carinii* and *P. wakefieldiae*: We aligned reads from *P. wakefieldiae*, *P. carinii, P. murina* and *P. jirovecii* to the *P. murina* reference genome (raw reads statistics are described in Supplementary Table 2). Our interspecies mapping strategy was designed to account for interspecies divergence of ~20% using LAST. We performed a clustering analysis of the SNP genotype data (total number of sites analyzed: 7,390,170; number of sites retained after filtering: 2,939,752). Figure 3a illustrates the distribution of each isolate according to distances. Hierarchical clustering cleanly separates samples from different species and indicates the majority of genetic variation within this data set consists of inter-species fixed differences. This allows computation of F_ST_ values comparing *P. murina* population (*n* = 12) to *P. carinii* (*n* = 7) and *P. wakefieldiae* (*n* = 5). F_ST_ genome scans reveal a significant population differentiation between *P. murina* and other species. In the comparison between *P. murina* and *P. wakefieldiae* populations, the genome-wide distribution of F_ST_ is unimodal with its highest peak at 0.99, which indicates that most of the *P. murina* genome is fully differentiated relative to *P. wakefieldiae*. In contrast, the comparison between *P. murina* and *P. carinii*, F_ST_ distribution is bimodal with two peaks at 0.9 and 0.99. A total of 86.3% and 93.7% of the 14,912 windows (5 kb) have an F_ST_ value > 0.8 in *P. murina*-*P. carinii* and *P. murina*-*P. wakefieldiae* comparisons, respectively. Genomic regions with F_ST_ < 0.5 represent 0.2% and 0.06%, respectively. Inspection of genomic regions with low differentiation (F_ST_ < 0.5) reveals a high incidence of SNPs, which suggests that low F_ST_ are caused by local increase in genetic diversity.

**Note 2:** Metabolic pathways.

Amino acids biosynthesis: The reduction of amino acid metabolism previously observed in *P. jirovecii*, *P. carinii* and *P. murina* (*24*) is also observed in *P. macacae*, *P. oryctolagi*, *P. canis* and *P. wakefieldiae* in this study. That is ~80% of the genes involved in amino acid biosynthesis are missing whereas most of their homologs are present in other Taphrinomycotina, which indicates that the loss of amino acid biosynthetic ability occurred in the LCA of *Pneumocystis*. All *Pneumocystis* have impaired capacity for assimilation of inorganic nitrogen and sulfur. Therefore, none of the 20 standard amino acids can be synthetized *de novo* although several can be derived from others. There is only one amino acid transporter (Ptr2) which is also conserved in all sequenced species. There is no amino acid permease in any sequenced *Pneumocystis* species. Transporters are significantly depleted.

*Pneumocystis* is defective in *de novo* biosynthesis of all standard amino acids and instead relies on scavenging from the host largely through mitochondrion and vacuolar associated amino-acid transporters. The unique amino acid transporter Avt3 is conserved in *P. jirovecii*, *P. macacae* and *P. oryctolagi*, but lost in the other 4 species sequenced. However, the directionality of Avt3 is unclear because it is usually used to import compounds from cytoplasm to help regulate intracellular levels (*28*). The polyamine transporter Aqr1, which is believed to be essential for *Pneumocystis* to scavenge polyamine (*29*), is only present in *P. jirovecii*, *P. macacae* and *P. canis*. Similarly, Aqr1 is an internal membrane protein involved in the excretion of excess of amino acids (*30*), thus it is unclear whether the Aqr1 transporter can also be used to import amino acids from the extracellular environment. These differences suggest that they are not essential to *Pneumocystis* survival and the loss may be due to a stochastic event. Apart from the few genes mentioned above, we found no significant differences in metabolic pathways among *Pneumocystis* species (Wilcoxon signed-rank test *p*-value = 0.7).

Serine/glycine biosynthesis: The serine/glycine biosynthesis pathway is probably nonfunctional in *Pneumocystis* because the key enzyme isocitrate lyase 1 as well as half of the 13 genes are absent in all *Pneumocystis*. The glycine cleavage system (glycine decarboxylase complex) which catalyzes the degradation of glycine is likely functional because nearly all species possess three copies of glycine decarboxylases (Gcv1, Gcv2, Gcv3). The only exception is *P. oryctolagi*, which lack two of three glycine decarboxylases and thus is probably unable to recycle glycine. *P. canis* genome lacks the glycine hydroxymethyltransferase SHM1, which is involved in the interconversion of serine and glycine.

Sulfur metabolism: Sulfur metabolism and synthesis of precursors for sulfur containing amino acids (methionine, cysteine, homocysteine, and taurine) is probably deficient because 13 of 22 genes of the pathway are lost in *Pneumocystis*. Key enzymes such as bifunctional cysteine synthase or cystathionine gamma-synthase are missing in all *Pneumocystis*. The cystathionine gamma lyase (Cys3), which is involved in the step 2 of synthesis of L-cysteine from L-homocysteine and L-serine, is missing in *P. oryctolagi* and *P. canis*. *P. oryctolagi* lacks the cystathionine beta-synthase (Cys4), which is involved in the step 1 of synthesis of L-cysteine from L-homocysteine and L-serine. These results suggest that these compounds are acquired from the host.

Carbohydrate metabolism: All necessary genes for uptake and catabolism of glucose via glycolysis and the tricarboxylic acid (TCA) cycle are present. Genes involved in the conversation of fructose and mannose to glucose, and the synthesize of glycogen and trehalose are present in all species. Key enzymes that convert galactose and sucrose are missing (galactokinase and beta-fructofuranosidase). Genes for galactose metabolism are conserved in other Taphrinomycotina which show that this pathway is lost specifically in the *Pneumocystis* branch. Additional losses include two enzymes for glyoxylation, one key enzyme for gluconeogenesis and all enzymes for pyruvate fermentation. These findings suggest that the reliance on glucose for energy production is a general feature for *Pneumocystis* species as suggested in our previous study (*25*).

Fatty acid metabolism: As observed before (*25*), most of the genes involved in the fatty acid beta oxidation are missing which further supports that hypothesis that fatty acids are not an energy source for *Pneumocystis*. Instead, *Pneumocystis* organisms rely on glycerol although the biosynthetic machinery of glycerol from glycerone-phosphate or monoacylglycerol is lost in all species. Glycerol uptake and export proteins Gup1 and Fps1 are conserved in all species except *P. macacae* and *P. canis* which have lost Fps1.Interestingly, *P. oryctolagi* and *P. canis* genomes have a single copy of *Dga1* (2-acylglycerol O-acyltransferase 2 [EC:2.3.1.22]), which is involved in glycerolipid synthesis. This gene is lost in other *Pneumocystis* species.

Pantothenate *de novo* biosynthesis: Pantothenate *de novo* biosynthesis is lost in all *Pneumocystis* species, whereas the ability to convert the pantothenate to CoA is conserved in all *Pneumocystis*. Given that the pantothenate specific transporter is also lost (Fen2), we have previously suggested that *Pneumocystis* uses the CoA transporter Leu5 to scavenge the pantothenate or its downstream metabolites from the hosts (*25*).

Sterol metabolism: Lipid metabolism in *Pneumocystis* is comparable among species. We have previously suggested that *P. jirovecii*, *P. carinii* and *P. murina* can synthetize fecosterol and episterol, but are unable to convert them to ergosterol because of the absence of two late stage enzymes Erg3 and Erg5 in the two latter species (*25*). . Moreover, both human and rodent *Pneumocystis* lost the key enzyme Dhcr24 (24-dehydrocholesterol reductase; EC 1.3.1.72) required for cholesterol biosynthesis. *P. macacae*, *P. oryctolagi* and *P. canis* genomes encode homologs of Erg3 and Dhcr7 whereas *P. wakefieldiae* lacks both of them. All species lack the key enzyme Dhcr24 required for cholesterol biosynthesis. This suggests that none of these species can synthesize cholesterol but presumably have the ability to scavenge it from the host though the host the genes involved in cholesterol acquisition have not been identified.

Cofactor metabolism: The riboflavin biosynthetic pathway is conserved in all *Pneumocystis* with the exception of *P. canis* which lacks RIB7 (2,5-diamino-6-ribosylamino-4(3H)-pyrimidinone 5'-phosphate reductase), a key enzyme required to catalyze an early step in riboflavin biosynthesis (*26*). The newly sequenced species also lack genes for the *de novo* synthesis of vitamins B1 and H. Genes encoding siderophores are also largely missing. The potential plasma membrane transporter for each of these cofactors is however highly conserved in all Taphrinomycotina including *Pneumocystis*. The NAD *de novo* synthesis pathway is severely truncated in *P. jirovecii* with loss of 6 out 9 genes. *P. macacae* also presents a reduction in this pathway, although less severe with 5/9 genes lost (the addition compared to *P. jirovecii* is Bna6, nicotinate-nucleotide diphosphorylase (carboxylating). In contrast *P. oryctolagi* has 8/9 genes (only the Bna2 (indoleamine 2,3-dioxygenase is missing)). Bna2 gene is also lost in *Schizosaccharomyces and Protomyces* while Bna6 is lost in *Schizosaccharomyces, Protomyces* and *Neolecta*. Bna2 catalyzes the first step of tryptophan catabolism in order to supply *de novo* nicotinamide adenine dinucleotide (NAD) via the kynurenine. Bn6 is involved in the catabolism of quinolinic acid. *P. canis*, *P. carinii*, *P. murina* and *P. wakefieldiae* have all the enzymes required for NAD *de novo* synthesis. The NAD salvage pathway including the nicotinic acid transporter is conserved in all these species. Some of these genes are lost in *Schizosaccharomyces* and *Protomyces* (Bna1 [HAD1 3-hydroxyanthranilate 3,4-dioxygenase], Bna2). This might indicate that the catabolism of tryptophan and quinolinic acid are not used by *P. jirovecii*, *P. macacae* and *P. oryctolagi*.

B6 metablism:Vitamin B6, which is an essential metabolite involved in defense against cellular oxidative stress, is synthesized by the DXP-independent pathway (*27*). The pathway involves two enzymes: Pdx1 (Pyridoxal 5'-phosphate synthase) and Pdx2. We found that Pdx1 is conserved in all except primate *Pneumocystis* species (*P. jirovecii* and *P. macacae*) whereas Pdx2 is conserved only in rodent-infecting species (*P. carinii*, *P. murina* and *P. wakefieldiae*). The B6 salvage pathway involves three enzymes: Bud16 (putative pyridoxal kinase), Pdx3 (pyridoxamine-phosphate oxidase) and Tpn1 (plasma membrane pyridoxine transporter). We identified orthologs of Bud16 in all *Pneumocystis* genomes and Pdx3 in all but *P. oryctolagi*, while Tpn1 is absent in all *Pneumocystis*. This uneven gene distribution pattern suggests that the *de novo* synthetic pathway for vitamin B6 may be functional only in rodent *Pneumocystis*, and that other *Pneumocystis* might have developed alternative strategies to generate B6, including scavenging from the host intermediate metabolites for B6 synthesis.

**Figure S1.** The maximum clade credibility tree of *Pneumocystis* summarized by TreeAnnotator and plotted against stratigraphy using the strap package in R. The internal nodes of the tree are indicated with circles, where the circles mark nodes with posterior probability: plain black > 0.95, grey > 0.75.

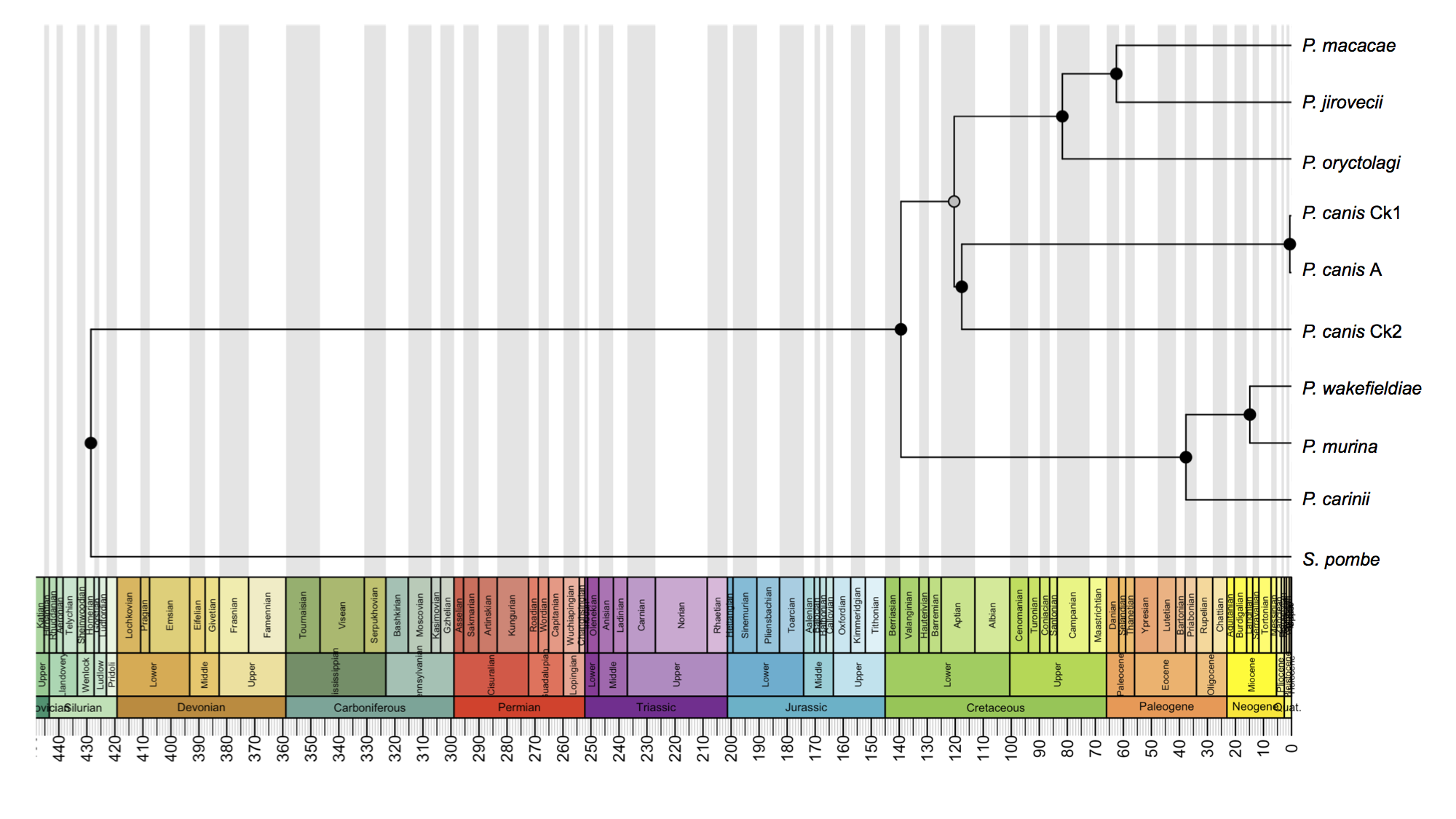

**Figure S2.** Maximum likelihood phylogeny constructed using a concatenated dataset of 15 protein coding genes from 33 *Pneumocystis* mitochondrial genomes including 7 *Pneumocystis* species. Pj, *P. jirovecii.* Pmac, *P.macacae*. Po, *P. oryctolagi*. Pcan, *P. canis*. Pcar, *P. carinii*. Pmur, *P. murina*. Pwk, *P. wakefieldiae*. The following genes were analyzed: cytochrome c oxidase subunits (*cox1*, *cox2, cox3*), ATP synthase F0 subunits (*atp6*, *atp8*, *atp9*), NADH dehydrogenase subunits (*nad1*, *nad2*, *nad3*, *nad5* and *nad6*), apocytochrome b (*cob*), ribonuclease P RNA (*rnpB*) and large and small subunit ribosomal RNAs (*rnl* and *rns*). Numbers in parentheses on the branches are the Shimodaira-Hasegawa [SH]-approximate likelihood ratio test (%) followed by the ultrafast bootstrap support (%).

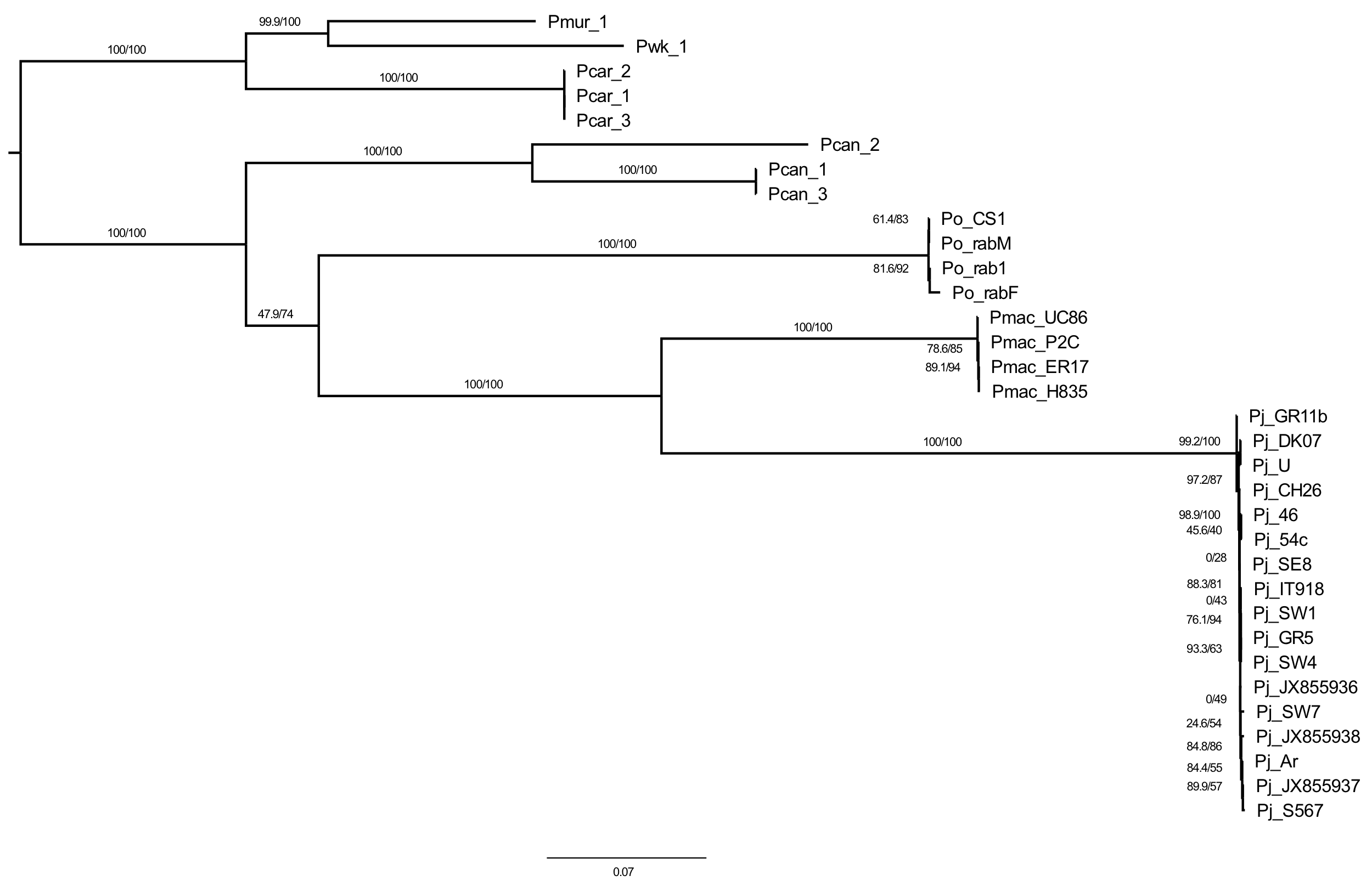

**Figure S3.** Genome-wide scans for footprints of natural selection in *Pneumocystis*. a, SNPs-based hierarchical clustering of *Pneumocystis* specimens (*n* = 59). b, Relative population divergence Fixation indexes (F_ST_) comparing *P. jirovecii* population to *P. macacae* and *P. oryctolagi*, respectively. c, Biplot of F_ST_ values comparing *P. jirovecii* population to *P. macacae* and *P. oryctolagi* populations colored according nucleotide diversities, showing that highly differentiated genomic regions tend to have a lower genetic diversity.

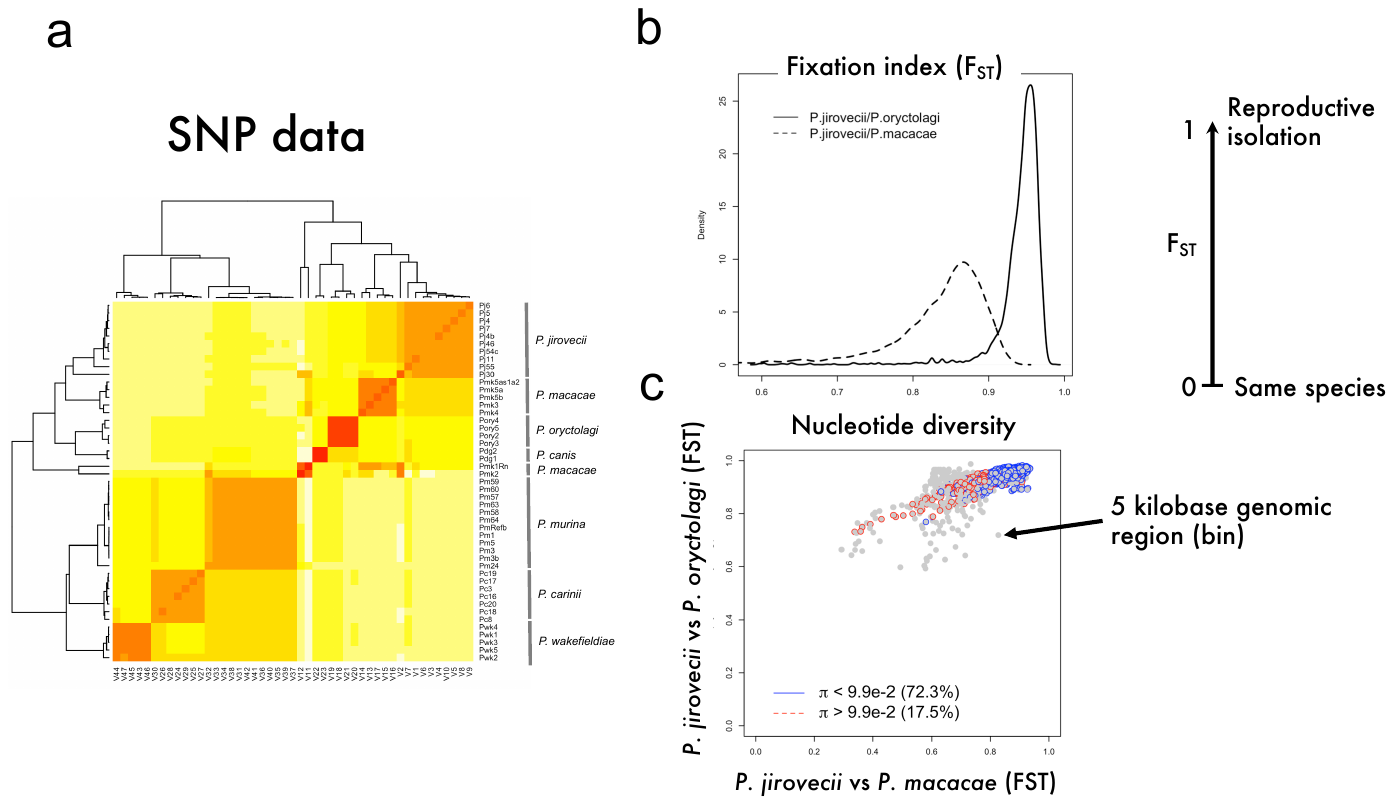

**Figure S4.** Evidence of ancient gene flow in rodent *Pneumocystis* only. Rooted phylogenetic network of seven *Pneumocystis* species as inferred by PhyloNet based on 1,718 one-to-one ortholog gene trees. Reticulations are shown as blue lines with inheritance probability of 64%.

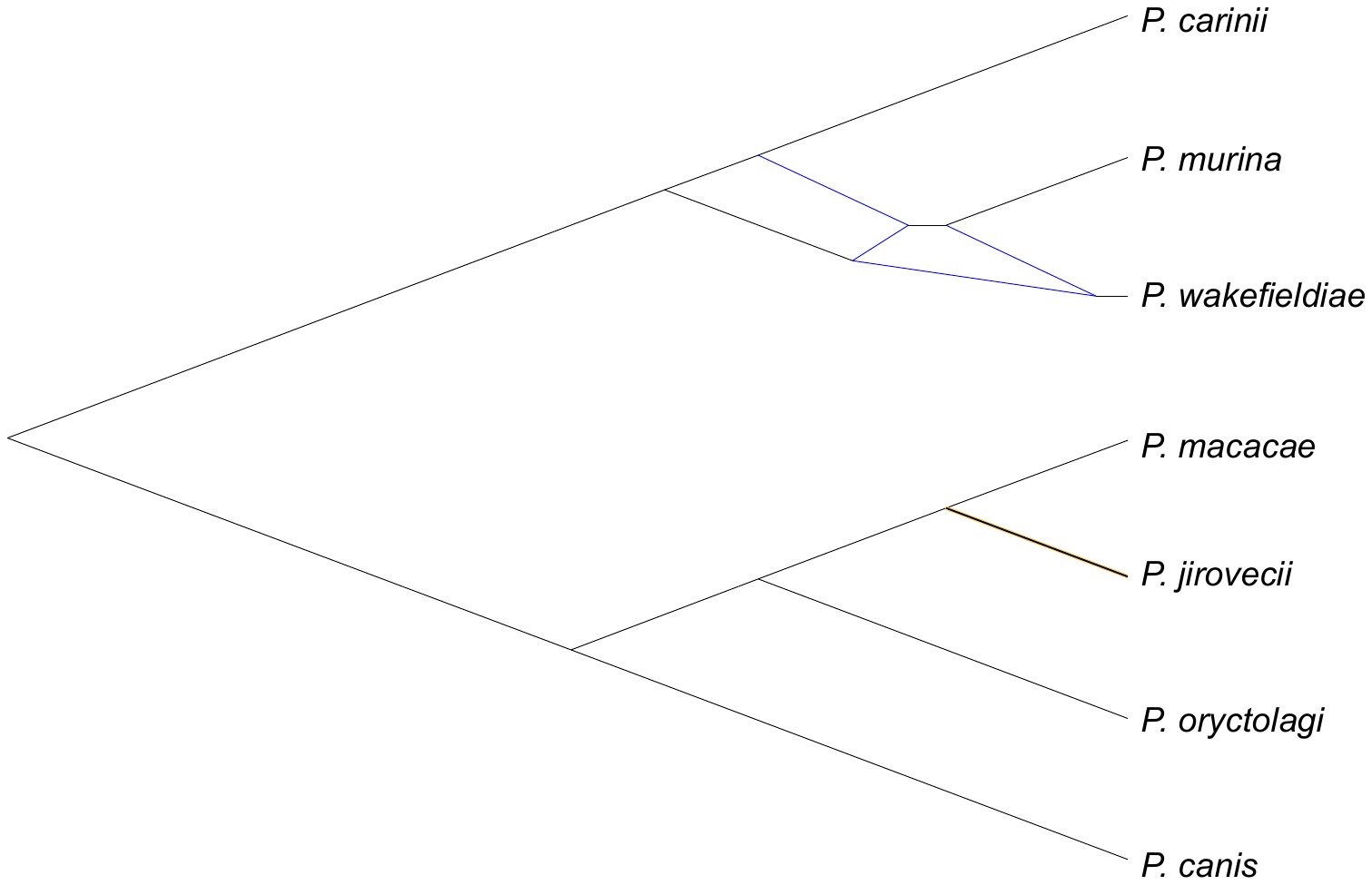

**Figure S5.** Evolution of arginine-glycine (RG) rich proteins in *P. macacae*. a, Sequence logos showing the frequency of amino acid composition in RG proteins. b, RNA-Seq of *P. macacae* genes showing high gene expression levels of RG genes relative to other genes*.* c, Analysis of a subset of *P. macacae* RGs (*n* = 6) using TOPALi based on the Difference of Sums of Squares method (DSS). Recombination is significantly detected at positions 75 to 150 of the alignment. Maximum likelihood of RG proteins. The blue star indicates the only protein with a signal peptide. d, Phylogenetic network of subtelomeric RG proteins showing reticulation events (possible recombination).

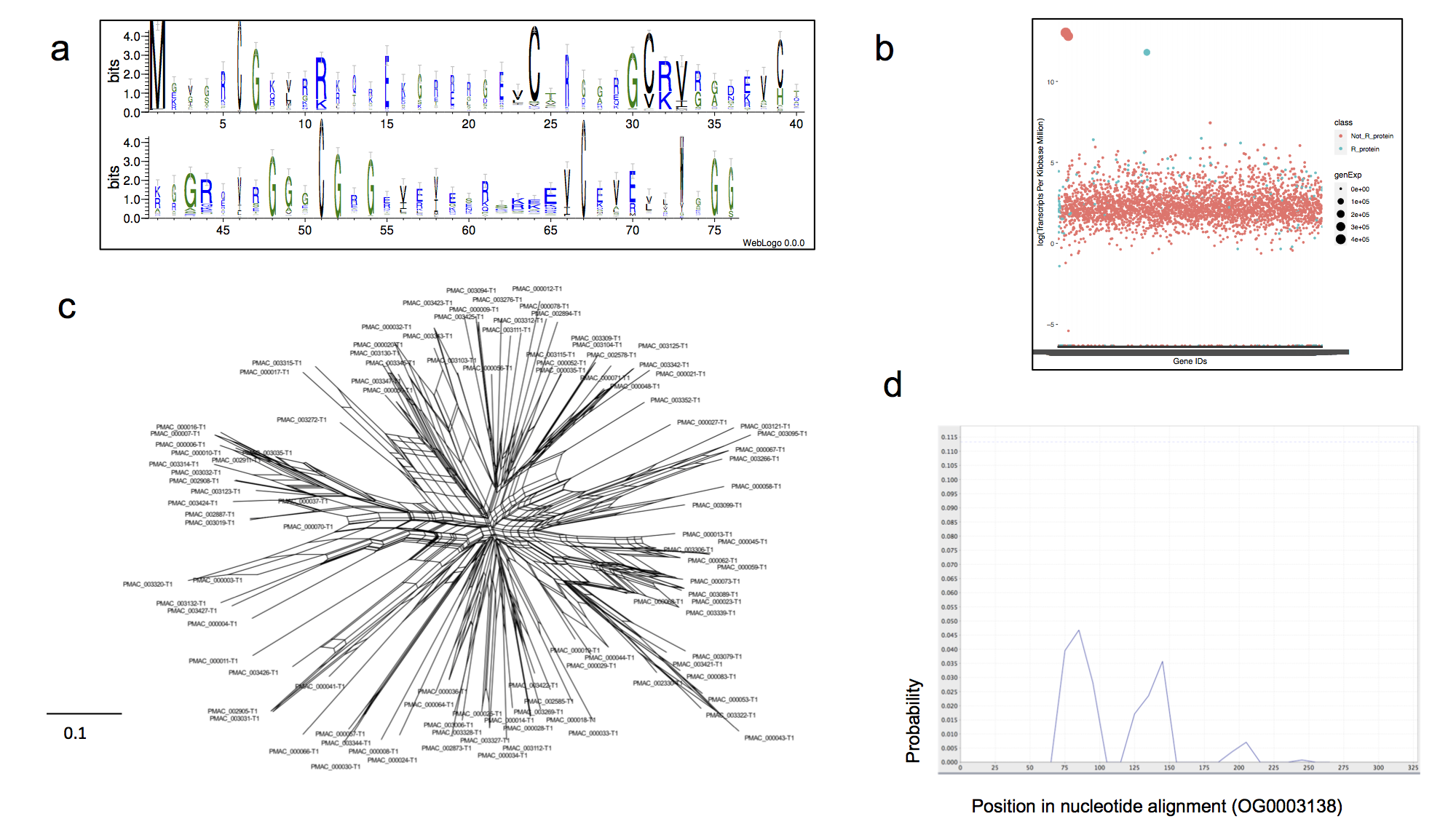

**Figure S6.** Heatmap showing gene family distribution in *Pneumocystis* species and related fungi.

Binary profiles of 7,720 gene families. Filled black squares denote presence of at least one member of the family in the genome or absence (empty) across the gene catalogs of seven *Pneumocystis* species, 11 related fungi from the Taphrinomycotina subphylum and five fungi from different subphyla. Taphrinomycotina fungi are presented in boxes at the bottom. TPOP, *Taphrina populina.* TW, *T. wiesneri*. TFLA, *T. flavoruba*. TDEF, *T. deformans*. SJAP, *Schizosaccharomyces japonicus*. SPOM, *S. pombe*. SCRYO, *Schizosaccharomyces cryophilus*. SOCT, *S. octosporus.* Prab, *Pneumocystis oryctolagi*. Pmac, *P. macacae.* PcanAus, *P. canis* A. PcanCk, *P. canis* Ck1. Pwk, *P. wakefieldiae*. PjRu7, *P. jirovecii*. PcB80, *P. carinii*. Pmur, *P. murina*. RHIOR, *Rhizopus* *oryzae*. Bd, *Batrachochytrium dendrobatidis.* NIRR, *Neolecta irregularis.* PROTO, *Protomyces inouyei.* CNEO, *Cryptococcus neoformans*. NCRA, *Neurospora crassa*. SAICO, *Saitoella complicata*.

**Figure S7.** Expansion of kexin peptidase families in *Pneumocystis*. Maximum likelihood phylogenetic tree of all kexin genes with hits to Pfam domain PF00082. A total of 13 kexin genes are present in *P. carinii* genome excluding those located in unassembled short contigs. All other species have only one copy except *P. wakefieldiae* which has three. The presence of a predicted GPI anchor and signal peptide is indicated by ‘+’. The number of predicted transmembrane regions is presented for each sequence. Sequences are labeled by a unique sequence identifier preceded by one of the species acronyms: PC, *P. carinii*; PWK, *P. wakefieldiae*; PM, *P. murina*; PCANCK, *P. canis* Ck1; PORYCT, *P. oryctolagi*; PJ, *P. jirovecii*; PMAC, *P. macacae*; SOCT, *Schizosaccharomyces octosporus*; Scryo, *S. cryophilus*; Spom, *S. pombe*; Sjap, *S. japonicus*.

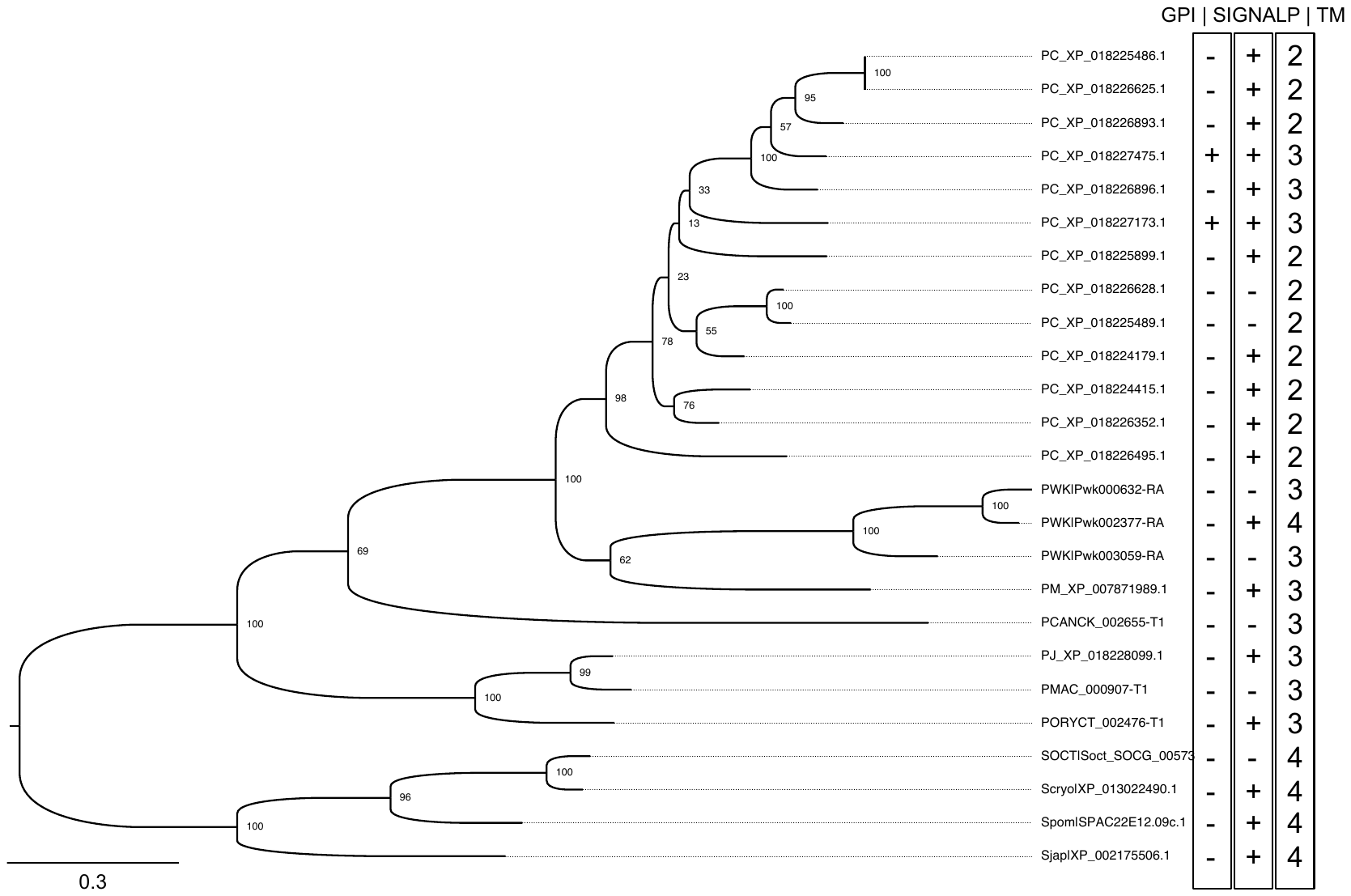

**Figure S8.** Evolutionary history of CFEM domains in *Pneumocystis* and other Taphrinomycotina. a, Maximum likelihood phylogeny of CFEM domain containing genes. b, Reconciliation of domain and /species trees showing CFEM domain evolutionary history. Sequences are labeled by a unique sequence identifier preceded by one of the species acronyms:

PJIR, *P. jirovecii*; PMAC, *P. macacae*; PORY, *P. oryctolagi*; PCAN, *P. canis* Ck1; PC, *P. carinii*; PMUR, *P. murina*; PWK, *P. wakefieldiae*. c, Sequence logo showing the frequency of amino acid composition in *Pneumocystis* CFEM domains.

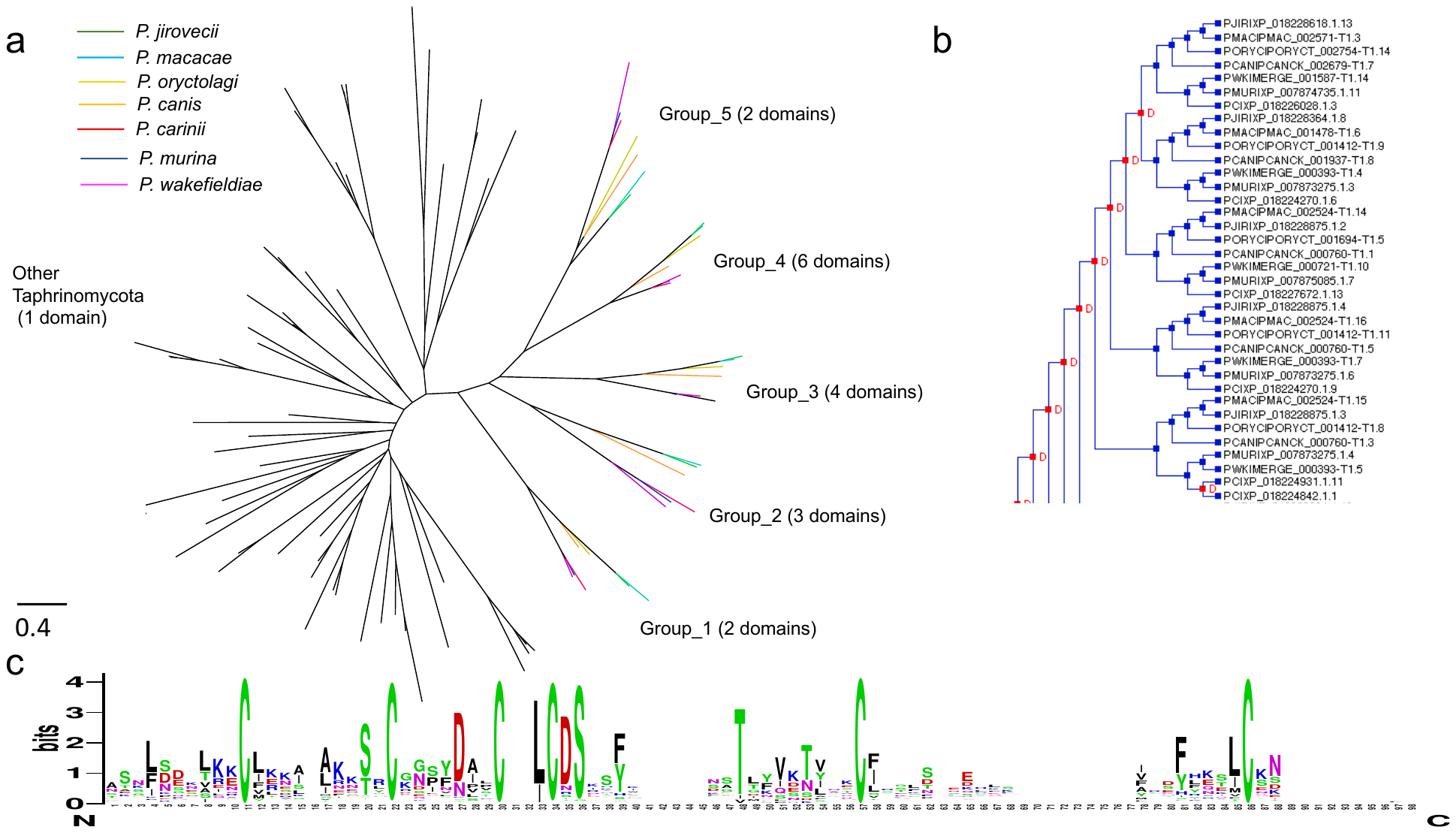

**Figure S9.** Evolutionary history of introns in *Pneumocystis* and Taphrinomycotina fungi. a) Graphical display of a representative protein alignment showing the gaps and intron position with respect to each of the aligned sequence. Nongap positions are indicated by grey boxes. Intron positions are indicated by small colored tags that have one, two, or three overlapping discs, indicating intron phase. Conserved (i.e., unambiguously aligned) intron-bearing sites are shown by dotted vertical lines. b) Bar chart of intron counts per species. c) Prediction of ancestral intron densities. The graphical display shows the inferred intron counts at the inner nodes and terminal taxa. The purple discs at the nodes have proportional diameters to the inferred densities. On each branch, the inferred number of intron losses (negative numbers) and gains (positive numbers) are presented. The orange and green bars have proportional heights to loss and gain amounts, respectively. Species are labelled with the following acronyms: *P. jirovecii* (pjir), *P. macacae* (pmac), *P. oryctolagi* (pory), *P. canis* (pcan), *P. carinii* (pcar), *P. murina* (pmur), *P. wakefieldiae* (pwk), *Schizosaccharomyces pombe* (spom), *Neolecta irregularis* (nirr), *Taphrina deformans* (tdef).

**Figure S10.** RAxML phylogeny and phylogenic networks of Msg genes. a, RAxML phylogeny of 482 Msg protein. b, Phylogenetic network of *P. jirovecii* and *P. macacae* *msg* genes only. Species are labelled as follows: *P. jirovecii* (Pj), *P. macacae* (Pmac), *P. carinii* (Pc), *P. murina* (Pm). c, Phylogenetic network of *P. carniii* and *P. murina and P. wakefieldiae*. Note that the full phylogenetic network was divided in two parts (b and c) to ease visualization. Complete phylogenetic network data is available at <https://github.com/ocisse/pneumocystis_evolution>.

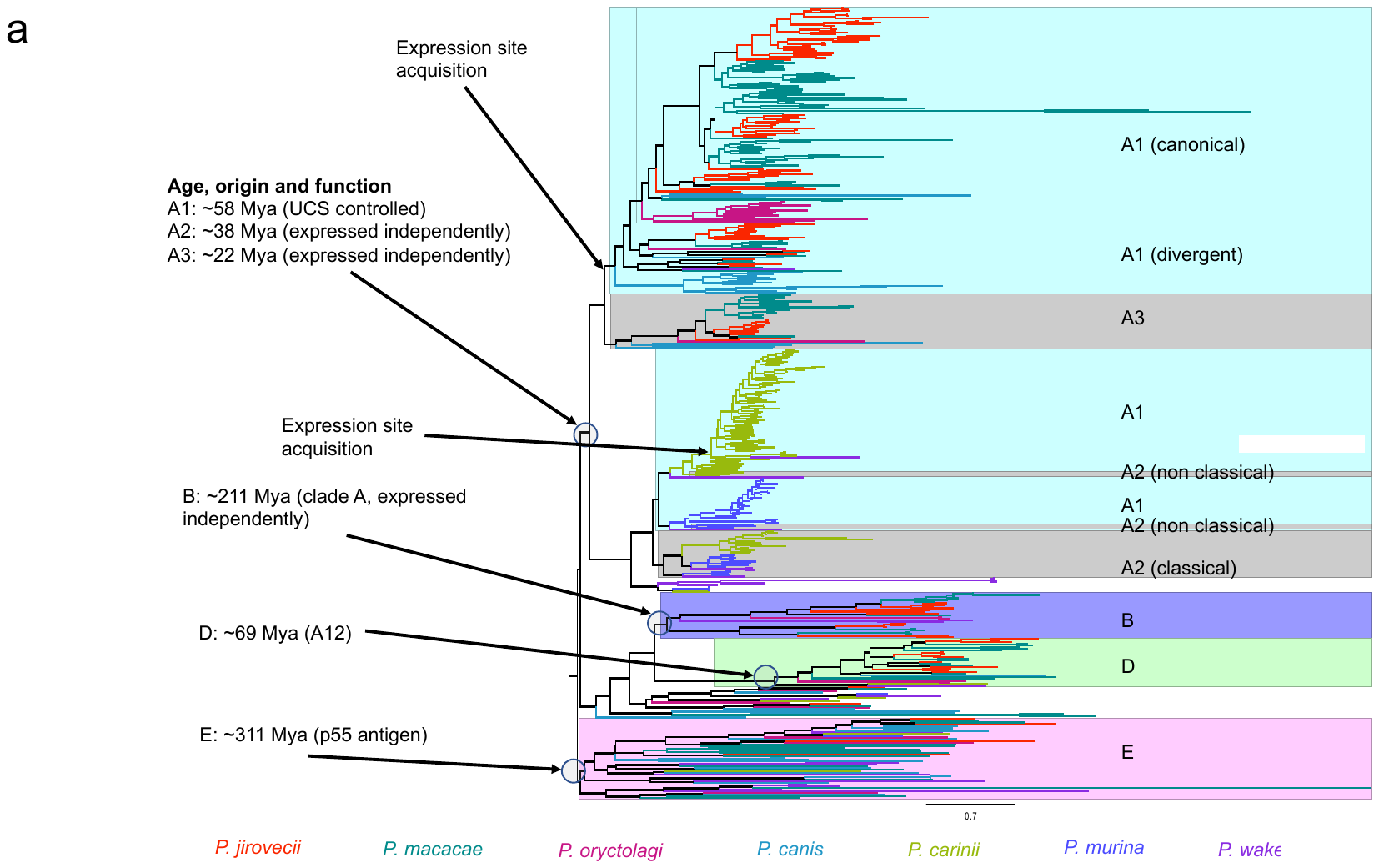

**Figure S11.** Phylodating of major surface glycoproteins in *Pneumocystis* and species divergence. Horizontal lines represent the 95% confidence intervals of Bayesian estimates and the dotted vertical line represents the emergence of the *Pneumocystis* genus.

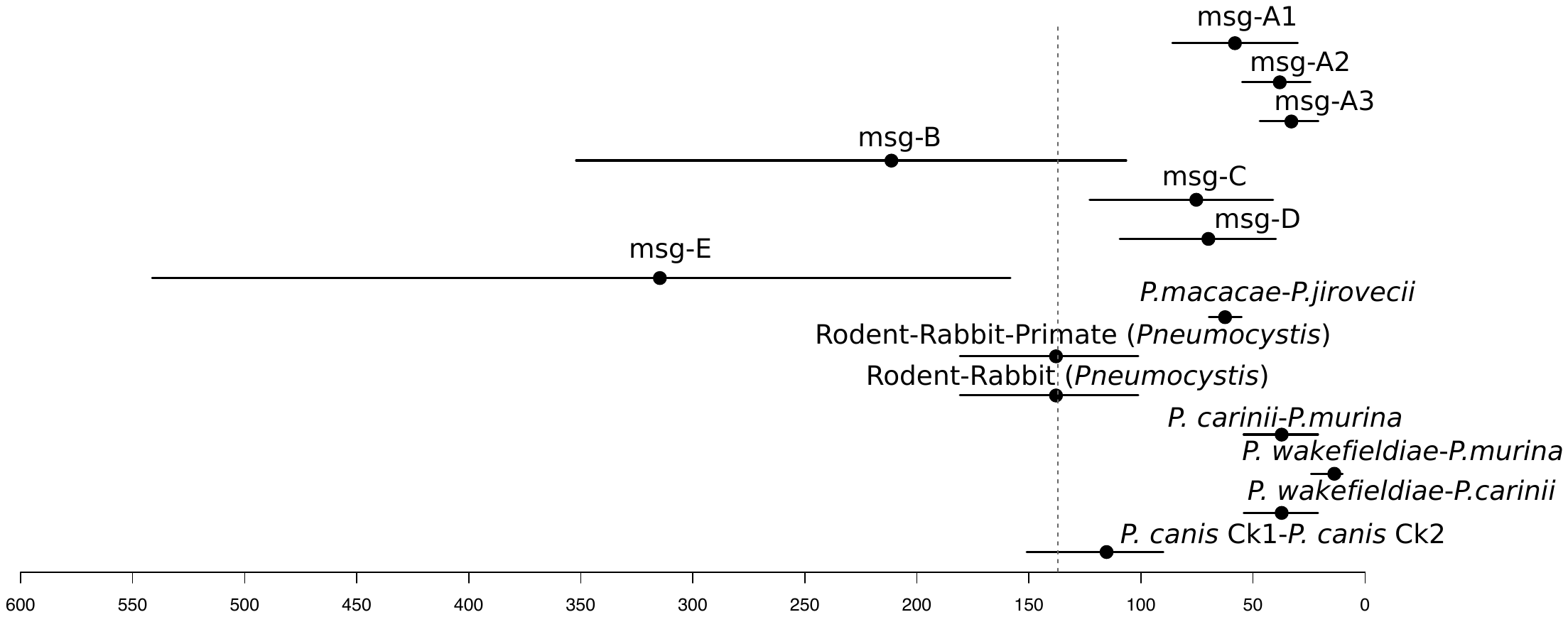

**Supplementary Table 1.** Clinical information and demographic data of individual samples used in this study *^a^*.

| **Species** | **Sample** | **Host species/strain** | **Collection dates (month, year)** | **Type of respiratory sample** | **Immune status/Condition** | **Host sex** | **Host age** | **Geographical location** |
| --- | --- | --- | --- | --- | --- | --- | --- | --- |
| ***P. jirovecii*** |  |  |  |  |  |  |  |  |
|  | Pj46 | *Homo sapiens* | n.a | BAL | n.a | n.a | n.a | Bethesda, Maryland, USA |
|  | Pj54c | *Homo sapiens* | 06/2018 | BAL | HIV infection | F | n.a | Bethesda, Maryland, USA |
|  | Pj55 | *Homo sapiens* | 11/2018 | BAL | Kidney transplantation | F | 27 | Chongqing, China |
| ***P. macacae*** |  |  |  |  |  |  |  |  |
|  | H835 | *Macaca mulatta* | 04/2013 | Autopsy lungs | Experimentally infected with SIV | n.a | n.a | Bethesda, Maryland, USA |
|  | P2C | *Macaca mulatta* | 06/2018 | Autopsy lungs | Experimentally infected with SIV | M | 6 years | Bethesda, Maryland, USA |
|  | CJ36 | *Macaca mulatta*/Indian | 06/2017 | Autopsy lungs | Experimentally infected with SIV | F | n.a | Tulane National Primate Research Center, Covington, LA |
|  | ER17 | *Macaca mulatta*/Indian | 12/2011 | FFPE sections | Experimentally infected with SIV | M | 8 years | Tulane National Primate Research Center, Covington, LA |
|  | UC86 | *Macaca mulatta*/Chinese | 02/1997 | FFPE sections | Experimentally infected with SIV | M | 1 year | CNPRC, Davis, CA |
|  | GL92 | *Macaca mulatta*/Indian | 01/2016 | FFPE sections | Experimentally infected with SIV | M | 9 years | Tulane National Primate Research Center, Covington, LA |
| ***P. oryctolagi*** |  |  |  |  |  |  |  |  |
|  | RABM | *Oryctolagus cuniculus*  /NZW/ Il2rg KO | 08/2017 | Lungs | SCID | M | 5 months | Ann Arbor, MI, Michigan |
|  | RABF | *Oryctolagus cuniculus* | n.a/2005 | Lungs | Spontaneous PCP at weaning | n.a | n.a | Lille, France |
|  | RAB1 | *Oryctolagus cuniculus* | n.a/2005 | Lungs | IBC | n.a | n.a | Lille, France |
|  | RAB2 | *Oryctolagus cuniculus* | n.a/2005 | Lungs | IBC | n.a | n.a | Lille, France |
| ***P. canis*** |  |  |  |  |  |  |  |  |
|  | Ck | *Canis lupus familiaris*  /Cavalier King Charles Spaniel | 10/1994 | FFPE sections | Suspected immunodeficiency | M | 1 year | Helsinki, Finland |
|  | A | *Canis lupus familiaris*  /Whippet mixed | 02/2013 | Lungs | n.a | M | 3 years | Vetmeduni, Austria |
| ***P. wakefieldiae/P. carinii*** |  |  |  |  |  |  |  |  |
|  | Pw2A | *Rattus norvegicus*/Long Evans rat | 09/1995 | Lungs | IBC | n.a | n.a | Cincinnati, OH, USA |
|  | PwC1 | *Rattus norvegicus*/Brown Norway rat | 10/2000 | Lungs | IBC | M | 13 weeks | Cincinnati, OH, USA |
|  | Pw1A | *Rattus norvegicus*/Brown Norway rat | 10/1996 | Lungs | IBC | n.a | n.a | Cincinnati, OH, USA |
|  | Pw3A | *Rattus norvegicus*/Brown Norway rat | 09/2002 | Lungs | IBC | M | 13 weeks | Cincinnati, OH, USA |
|  | PwRN | *Rattus norvegicus* | 10/1992 | Lungs | IBC | n.a | n.a | Cincinnati, OH, USA |
| ***P. murina*** |  |  |  |  |  |  |  |  |
|  | D3A | CD40 L KO mice/A549 cells | 07/2018 | Lungs | n.a. | n.a | 3 days | Bethesda, Maryland, USA |

Abbreviations: BAL, bronchoalveolar lavage; FFPE, formalin-fixed paraffin-embedded tissues; IBC, immunosuppressed by corticosteroid administration; PCP, *Pneumocystis* Pneumonia; NZW/Il2rg KO, Interleukin-2 receptor-ɣ knock out mutant rabbits; SCID, severe combined immunodeficiency; SIV, Simian immunodeficiency virus;

n.a., not available.

*^a^* Public raw sequencing data used in this study are presented separately in Supplementary Table 2.

**Supplementary Table 2.** Statistics and a posteriori classification of reads used in this study.

| Species | *n* | Sample ID | Type | Host ID | Raw paired reads (millions) | *Pneumocystis* reads (%) | Median coverage | Data source | NCBI BioProject/SRA accession # |
| --- | --- | --- | --- | --- | --- | --- | --- | --- | --- |
| *P. jirovecii* |  |  |  |  |  |  |  |  |  |
|  | 1 | Pj46 | DNA | 46 | 26.6 | 0.1 | 4.8 | This study | SRR12304674 |
|  | 2 | Pj54c | DNA | 54c | 46.2 | 0.8 | 47.4 | This study | SRR12304306 |
|  | 3 | Pj55 | DNA | 55 | 24.6 | 6.8 | 10.1 | This study | SRR12300271 |
|  | 4 | RU817b | DNA | RU | 10.7 | 65.0 | 85.4 | This study | n.a. |
|  | 5 | RU7 | DNA | RU | 70.5 | 47.0 | 342.3 | Ma *et al.* ^(^*^25^*^)^ | SRR1043749 |
|  | 6 | RU12 | DNA | RU | 70.6 | 48.0 | 353.3 | Ma *et al.* ^(^*^25^*^)^ | SRR1043747 |
|  | 7 | RU817 | DNA | RU | 70.6 | 46.0 | 338.7 | Ma *et al.* ^(^*^25^*^)^ | SRR1043750 |
|  | 8 | SE8 | DNA | SE8 | 95.5 | 15.0 | 1,364.5 | Cissé *et al.* ^(^*^10^*^)^ | PRJEB2702 |
|  | 9 | Z | DNA | Z | 89.6 | 9.4 | 11.6 | Ma *et al.* ^(^*^25^*^)^ | SRR5822089,  SRR5825481,  SRR5825487 |
|  | 10 | W | DNA | W | 91.8 | 17.0 | 64.6 | Ma e*t al.* ^(^*^25^*^)^ | SRR5822088 |
|  | 11 | RN1 | RNA | RN1 | 21.6 | 5.3 | 11.8 | Cissé *et al.* ^(^*^10^*^)^ | PRJEB3063 |
|  | 12 | RN2 | RNA | RN2 | 25.5 | 0.6 | 6.7 | Cissé *et al.* ^(^*^27^*^)^ | SRR5822079 |
| *P. macacae* |  |  |  |  |  |  |  |  |  |
|  | 1 | M2 | DNA | H835 | 7.7 | 1.1 | 4.6 | Cissé *et al.* ^(^*^27^*^)^ | SRR12323870 |
|  | 2 | MP3 | DNA | H835 | 3.7 | 0.3 | 1.5 | Cissé *et al.* ^(^*^27^*^)^ | SRR12323870 |
|  | 3 | H835 | DNA | H835 | 0.6 | 1.0 | 1.9 | Cissé *et al.* ^(^*^27^*^)^ | SRR12323870 |
|  | 5 | CJ36 | DNA | CJ36 | 35.4 | 0.0 | 5.4 | This study | SRR12323878 |
|  | 6 | ER17 | DNA | ER17 | 52.2 | 0.5 | 8.7 | This study | PRJNA648108 |
|  | 7 | UC86 | DNA | UC86 | 38.0 | 4.4 | 8.1 | This study | PRJNA648112 |
|  | 8 | P2C | DNA | P2C | 61.7 | 68.2 | 1,394.4 | This study | SRR11785744 |
|  | 9 | S1A2 | DNA | P2C | 24.0 | 19.3 | 151.8 | This study | SRR11785744 |
|  | 10 | P2C | DNA | P2C | 1.6*^b^* | 5.0 | n.a | This study | SRR11785744 |
|  | 10 | GL92 | DNA | GL92 | 29.8 | 2.1 | 4.5 | This study | PRJNA648115 |
|  | 11 | RN1 | RNA | H835 | 18.2 | 1.9 | 13.4 | Cissé et al. 2018 ^(^*^27^*^)^ | SRR5822119 |
|  | 12 | RN2 | RNA | P2C | 22.0 | 91.3 | 12.0 | This study | SRR11785742 |
| *P. oryctolagi* |  |  |  |  |  |  |  |  |  |
|  | 1 | RABF | DNA | RABF | 73.0 | 3.6 | 9.6 | This study | SRR11789048 |
|  | 2 | RABM | DNA | RABM | 73.5 | 4.4 | 11.1 | This study | [SRR11789047](https://dataview.ncbi.nlm.nih.gov/object/SRR11789047) |
|  | 3 | RAB1 | DNA | RAB1 | 17.7 | 1.0 | 9.2 | This study | SRR11789046 |
|  | 4 | RAB2 | DNA | RAB2 | 20.7 | 1.1 | 25.4 | This study | SRR11789045 |
| *P. canis* |  |  |  |  |  |  |  |  |  |
|  | 1 | Ck | DNA | Ck | 111.4 | 1.9 | 17.1 | This study | SRR11795436 |
|  | 2 | A | DNA | A | 38.8 | 5.1 | 15.3 | This study | SRR11908556 |
| *P. wakefieldiae* |  |  |  |  |  |  |  |  |  |
|  | 1 | 2A | DNA | Pw2A | 23.2 | 17.6 | 167.4 | This study | SRR11794283 |
|  | 2 | C1 | DNA | PwC1 | 29.5 | 0.5 | 2.0 | This study | SRR11794286 |
|  | 3 | 1A | DNA | Pw1A | 67.5 | 79.7 | 2,193.2 | This study | SRR11794285 |
|  | 4 | 3A | DNA | Pw3A | 66.3 | 33.2 | 899.2 | This study | SRR11794284 |
|  | 5 | 2A | RNA | RNA | 39.7 | 25.2 | n.a | This study | SRR11794282 |
| *P. carinii* |  |  |  |  |  |  |  |  |  |
|  | 1 | B80 | DNA | B80 | 28.8 | 56.8 | 389.6 | Ma *et al.* ^(^*^25^*^)^ | SRR1043726, SRR1043727 |
|  | 2 | B50 | DNA | B50 | 11.7 | 41.4 | 116.8 | Ma *et al.* ^(^*^25^*^)^ | SRR1043724 |
|  | 3 | SE6 | DNA | SE6 | 28.6 | 21.3 | 53.6 | Cissé *et al.* ^(^*^10^*^)^ | ERR047567 |
|  | 4 | Pc1 | RNA | 70954 | 14.3 | 4.9 | 160.0 | Ma *et al.* ^(^*^25^*^)^ | SRR2156857 |
|  | 5 | Pc2 | RNA | 70955 | 13.7 | 6.3 | 188.4 | Ma *et al.* ^(^*^25^*^)^ | SRR2156986 |
|  | 6 | Pc3 | RNA | 70956 | 13.5 | 5.6 | 162.2 | Ma *et al.* ^(^*^25^*^)^ | SRR2156987 |
|  | 7 | 2A | DNA | Pc2A | 23.2 | 0.7 | 6.4 | This study | SRR11794283 |
|  | 8 | C1 | DNA | PcC1 | 29.5 | 7.5 | 87.1 | This study | SRR11794286 |
|  | 9 | 1A | DNA | Pc1A | 67.5 | 0.4 | 10.9 | This study | SRR11794285 |
|  | 10 | 3A | DNA | Pc3A | 66.3 | 3.8 | 98.4 | This study | SRR11794284 |
|  | 11 | 2A | RNA | Pc2A | 39.7 | 26.1 | n.a | This study | SRR11794282 |
| *P. murina* |  |  |  |  |  |  |  |  |  |
|  | 1 | Pm1 | DNA | C1 | 17.6 | 23.8 | 107.1 | Ma *et al.* ^(^*^25^*^)^ | SRR6001186 |
|  | 2 | Da3 | DNA | Da3 | 23.2 | 4.8 | 24.1 | Ma *et al.* ^(^*^25^*^)^ | SRR6001196 |
|  | 3 | Da1 | DNA | Da1 | 19.3 | 3.8 | 15.2 | Ma *et al.* ^(^*^25^*^)^ | SRR6001188 |
|  | 4 | MS96 | DNA | MS96 | 19.9 | 9.8 | 46.5 | Ma *et al.* ^(^*^25^*^)^ | SRR6001185 |
|  | 5 | A123 | DNA | A123 | 18.9 | 29.2 | 139.9 | Ma *et al.* ^(^*^25^*^)^ | SRR6001197 |
|  | 6 | Pm24 | RNA | A549_PC | 20.8 | 4.8 | 27.5 | This study | SRR12313714 |
|  | 7 | Pm57 | RNA | Pm57 | 48.9*^a^* | 19.2 | 4.2 | Cushion *et al.* ^(^*^28^*^)^ | SRR6721590 |
|  | 8 | Pm58 | RNA | Pm58 | 43.6 *^a^* | 18.3 | 5.8 | Cushion *et al.* ^(^*^28^*^)^ | SRR6721591 |
|  | 9 | Pm59 | RNA | Pm59 | 49.1 *^a^* | 19.4 | 3.8 | Cushion *et al.* ^(^*^28^*^)^ | SRR6721592 |
|  | 10 | Pm60 | RNA | Pm60 | 42.3 *^a^* | 14.4 | 4.6 | Cushion *et al.* ^(^*^28^*^)^ | SRR6721593 |
|  | 11 | Pm63 | RNA | Pm63 | 44.3 *^a^* | 11.6 | 4.6 | Cushion *et al.* ^(^*^28^*^)^ | SRR6721596 |
|  | 12 | Pm64 | RNA | Pm64 | 46.6 *^a^* | 19.1 | 5.6 | Cushion *et al.* ^(^*^28^*^)^ | SRR6721597 |

**Supplementary Table 3.** Statistics of different *Pneumocystis* genome assemblies.

|  |  | **Species** | | | | | | | | |
| --- | --- | --- | --- | --- | --- | --- | --- | --- | --- | --- |
|  | **Features** | ***P. macacae*** | ***P. oryctolagi*** | ***P. canis* Ck1** | ***P. canis* Ck2** | ***P. canis* A** | ***P. wakefieldiae*** | ***P. jirovecii*** | ***P. carinii*** | ***P. murina*** |
| **Genomic DNA** |  |  |  |  |  |  |  |  |  |  |
|  | Source | This study | This study | This study | This study | This study | This study | Ma et al. ^(^*^25^*^)^ | Ma *et al.* ^(^*^25^*^)^ | Ma *et al.* ^(^*^25^*^)^ |
|  | Host | Macaque | Rabbit | Dog Ck | Dog Ck | Dog A | Rat | Human | Rat | Mouse |
|  | Strain | P2C | CS1 | Ck1*^a^* | Ck2 *^a^* | A | 2A | RU7 | B80 | B123 |
|  | DNA amplification | None | WGA | None | None | None | None | None | None | None |
| **Scaffolds** |  |  |  |  |  |  |  |  |  |  |
|  | Sequencing technology | Nanopore + Illumina | Illumina | Illumina | Illumina | Illumina | Illumina | PacBio + Illumina | PacBio + Illumina | PacBio + Illumina |
|  | Total length (Mb) | 8.2 | 7.6 | 7.9 | 3.6 | 7.4 | 7.3 | 8.3 | 7.6 | 7.4 |
|  | Number | 16*^b^* | 38 | 78 | 315 | 33 | 17 | 20*^b^* | 17*^b^* | 17*^b^* |
|  | Coverage | 1,394.4 | 13.8 | 17.2 | n.a | 15.3 | 2663.2 | 342.3 | 389.6 | 139.9 |
|  | N50 (kb) | 505.3 | 534.6 | 457.1 | 174.4 | 422.4 | 480.6 | 454.5 | 465.1 | 491.3 |
|  | GC content (%) | 29.1 | 28.6 | 26.3 | 29.7 | 25.9 | 29.8 | 28.7 | 27.8 | 26.9 |
|  | With telomeric motif (*n*) | 7 | 2 | 2 | 4 | 0 | 5 | 7 | 6 | 9 |
|  | Karyotype (*n*) | n.a | 14 ^(^*^29^*^)^ | n.a | n.a | n.a | 14 ^(^*^11^*^)^ | 17-19 ^(^*^30^*^)^ | 13-15 ^(^*^31^*^)^ | 17 ^(^*^32^*^)^ |
| **Nuclear genome assembly completeness (%)** |  |  |  |  |  |  |  |  |  |  |
|  | CEGMA (%) | 89.9 | 89.9 | 91.1 | 14.9 | 93.1 | 93.1 | 91.1 | 91.3 | 91.9 |
|  | BUSCO (%) | 86.2 | 92.4 | 90.7 | 15.6 | 91.8 | 92.4 | 95.1 | 94.8 | 94.4 |
|  | FGMP (%) | 92.2 | 89.6 | 93.4 | 49.2 | 93.4 | 92.6 | 93.1 | 94.4 | 92.2 |
| **Nuclear annotation statistics** |  |  |  |  |  |  |  |  |  |  |
|  | Protein-coding genes (*n*)*^c^* | 3,427 | 2,961 | 3,476 | 2,135 | 3,077 | 3,221 | 3,765 | 3,646 | 3,838 |
|  | Ribosomal RNA genes (*n*) | 5 | 10 | 5 | 5 | 5 | 5 | 5 | 5 | 5 |
|  | Transfer RNA genes (*n*) | 47 | 47 | 46 | 18 | 47 | 47 | 46 | 45 | 47 |
|  | Exons (*n*) | 21,608 | 21,574 | 21,632 | 8,544 | 20,216 | 23,126 | 21,770 | 21,808 | 22,085 |
|  | Exons per gene (*n*) | 6 | 6 | 6 | 5 | 6 | 7 | 6 | 6 | 6 |
|  | Orthogroups (*n*) | 3,280 | 3,102 | 3,395 | 1,296 | 3,060 | 3,133 | 3,643 | 3,602 | 3,574 |
|  | Orphans (*n*) | 190 | 204 | 80 | 44 | 17 | 25 | 120 | 42 | 50 |
|  | Msg (*n*) | 106 | 9*^c^* | 22 | 36 *^c^* | 23 | 17 | 161 | 93 | 61 |
|  | Repeats (%) *^d^* | 2.6 | 2.8 | 2.7 | 1.6 | 2.7 | 1.8 | 2.7 | 3.0 | 2.4 |
| **Mitochondrial genome statistics** |  |  |  |  |  |  |  |  |  |  |
|  | Source | This study | This study | This study | This study | This study | This study | Ma *et al.* ^(^*^33^*^)^ | Ma *et al.*^(^*^33^*^)^ | Ma *et al.* ^(^*^33^*^)^ |
|  | Total length (bp) | 21,232 | 24,512 | 21,750 | 21,413 | 21,368 | 23,752 | 35,626 | 26,119 | 24,608 |
|  | Scaffolds or contigs (*n*) | 1 | 1 | 1 | 1 | 1 | 1 | 1 | 1 | 1 |
|  | Predicted topology | Circular | Linear | Linear | Linear | Linear | Linear | Circular | Linear | Linear |
|  | GC content (%) | 28.9 | 31.5 | 29.3 | 29.4 | 29.4 | 30.0 | 25.7 | 29.8 | 29.8 |
|  | Protein-coding genes (*n*)*^e^* | 15 | 15 | 15 | 15 | 15 | 15 | 15 | 15 | 15 |
|  | rRNA genes (*n*) | 3 | 3 | 3 | 3 | 3 | 3 | 3 | 3 | 3 |
|  | tRNA genes (*n*) | 22 | 22 | 22 | 23 | 23 | 22 | 25 | 25 | 28 |

*^a^ P. canis* genome assemblies Ck1 and Ck2 are from the same dog Ck.

*^b^* Excluding short contigs for *msg* genes.

*^c^* Possibly incomplete due to difficulties in assembling full-length *msg* genes from Illumina reads alone.

*^d^* DNA transposons, retrotransposons and simple low complexity AT rich repeats.

*^e^* Including one hypothetic protein-coding gene (orf195).

Abbreviations: WGA: Whole Genome Amplification; CEGMA, Core Eukaryotic Genes Mapping Approach; BUSCO, Benchmarking Universal Single-Copy Orthologs; FGMP, Fungal Genome Mapping Pipeline; n.a., not available.

**Supplementary Table 4.** Genome rearrangements among different *Pneumocystis* species.

| GR | GO | Syntenic blocks (*n*)*^a^* | Inversions (*n*)*^b^* | Genome rearrangement breakpoints*^c^* | | | | | | |
| --- | --- | --- | --- | --- | --- | --- | --- | --- | --- | --- |
|  |  |  |  | Total refined (*n*) | Genic (*n*) | IGS | Repeats*^d^* | Divergence (%) | Telomeric | Msg |
| *P. jirovecii* | *P. macacae* | 43 | 23 | 29 | 13 | 16 | 19 | 15.7 | 2 | 0 |
| *P. macacae* | *P. jirovecii* | 43 | 23 | 33 | 25 | 8 | 11 |  | 0 | 0 |
| *P. jirovecii* | *P. oryctolagi* | 225 | 26 | 142 | 97 | 45 | 31 | 20.5 | 1 | 0 |
| *P. oryctolagi* | *P. jirovecii* | 225 | 26 | 132 | 84 | 48 | 24 |  | 4 | 0 |
| *P. jirovecii* | *P. canis* Ck1 | 168 | 48 | 82 | 42 | 40 | 31 | 21.3 | 2 | 0 |
| *P. canis* Ck1 | *P. jirovecii* | 168 | 48 | 79 | 40 | 39 | 20 |  | 1 | 0 |
| *P. jirovecii* | *P. carinii* | 134 | 53 | 82 | 44 | 38 | 34 | 22.6 | 2 | 0 |
| *P. carinii* | *P. jirovecii* | 134 | 53 | 85 | 70 | 15 | 19 |  | 2 | 0 |
| *P. jirovecii* | *P. murina* | 112 | 45 | 79 | 39 | 40 | 35 | 22.4 | 2 | 0 |
| *P. murina* | *P. jirovecii* | 112 | 45 | 82 | 60 | 21 | 10 |  | 0 | 0 |
| *P. jirovecii* | *P. wakefieldiae* | 128 | 46 | 86 | 46 | 40 | 38 | 22.9 | 1 | 0 |
| *P.wakefieldiae* | *P. jirovecii* | 128 | 46 | 89 | 58 | 31 | 10 |  | 0 | 0 |
| *P.wakefieldiae* | *P. murina* | 29 | 9 | 10 | 0 | 10 | 1 | 12.0 | 0 | 0 |
| *P. murina* | *P. wakefieldiae* | 29 | 9 | 10 | 5 | 5 | 1 |  | 0 | 0 |
| *P.wakefieldiae* | *P. carinii* | 44 | 14 | 20 | 13 | 7 | 4 | 14.9 | 1 | 0 |
| *P. carinii* | *P. wakefieldiae* | 44 | 14 | 20 | 13 | 7 | 4 |  | 1 | 0 |
| *P. carinii* | *P. murina* | 29 | 14 | 10 | 3 | 7 | 2 | 12.3 | 0 | 0 |
| *P. murina* | *P. carinii* | 29 | 14 | 10 | 0 | 10 | 0 |  | 0 | 0 |
| *S.octoporus* | *S.cryophilus* | 71 | 19 | 30 | 18 | 12 | 1 | 20.9 | 0 | NA |
| *S.cryophilus* | *S.octoporus* | 71 | 19 | 24 | 17 | 7 | 2 |  | 0 | NA |
| *S. pombe* | *S.japonicus* | 708 | NA | 160 | 159 | 1 | 8 | 21.0 | 0 | NA |
| *S.japonicus* | *S. pombe* | 708 | NA | 160 | 154 | 6 | 0 |  | 0 | NA |

Abbreviations: BPs: breakpoints; IGS: intergenic spaces; GR: reference genome; GO: queried genome; Msg: major surface glycoprotein encoding regions.

*^a^* Syntenic blocks defined by MAUVE/GRIMM.

*^b^* Inversions were inferred using MAUVE.

*^c^* BPs were refined using Cassis and identified in the GR (reference genome) not in the GO.

*^d^* Repeats as identified using RepeatMasker. These elements might overlap with genic and IGS regions.

**Supplementary Table 5.** Pairwise nucleotide divergence (%) among *Pneumocystis* genomes.

| Species (Specimen) | *P. jirovecii* (RU7) | *P. jirovecii* (SE8) | *P. jirovecii*  (SE2178) | *P. macacae* (P2C) | *P. oryctolagi* (CS1) | *P. canis* (Ck1) | *P. canis* (Ck2) | *P. canis* (A) | *P carinii* (B80) | *P carinii* (Ccin) | *P. carinii* (SE6) | *P. murina* (B123) | *P. wakefieldiae* (2A) | Data source | NCBI accession # |
| --- | --- | --- | --- | --- | --- | --- | --- | --- | --- | --- | --- | --- | --- | --- | --- |
| P. jirovecii (RU7) | 0.0 |  |  |  |  |  |  |  |  |  |  |  |  | Ma et al. ^(^*^25^*^)^ | GCA_001477535.1 |
| P. jirovecii (SE8) | 0.2 | 0.0 |  |  |  |  |  |  |  |  |  |  |  | Cisse et al.^(^*^10^*^)^ | GCA_000333975.2 |
| P. jirovecii (SE2178) | 0.8 | 0.5 | 0.0 |  |  |  |  |  |  |  |  |  |  | Schmid et al. ^(^*^34^*^)^ | GCA_002571455.1 |
| P. macacae (P2C) | 15.7 | 15.1 | 15.6 | 0.0 |  |  |  |  |  |  |  |  |  | This study | JABMLO000000000 |
| P. oryctolagi (CS1) | 20.5 | 20.4 | 20.5 | 20.8 | 0.0 |  |  |  |  |  |  |  |  | This study | JABTEG000000000 |
| P. canis (Ck1) | 21.3 | 21.2 | 21.2 | 21.5 | 21.6 | 0.0 |  |  |  |  |  |  |  | This study | JABTEF000000000 |
| P. canis (Ck2) | 20.6 | 20.6 | 20.6 | 20.7 | 20.7 | 15.5 | 0.0 |  |  |  |  |  |  | This study | JABVCJ000000000 |
| P. canis (A) | 21.3 | 21.2 | 21.3 | 21.5 | 21.6 | 0.5 | 15.5 | 0.0 |  |  |  |  |  | This study | JACEFK000000000 |
| P. carinii (B80) | 22.6 | 22.5 | 22.6 | 22.8 | 22.8 | 22.5 | 21.0 | 22.5 | 0.0 |  |  |  |  | Ma et al. ^(^*^25^*^)^ | GCA_001477545.1 |
| P. carinii (Ccin) | 23.1 | 23.0 | 23.1 | 22.8 | 23.2 | 23.1 | 21.0 | 23.1 | 5.0 | 0.0 |  |  |  | Slaven et al. ^(^*^9^*^)^ | This study |
| P. carinii (SE6) | 21.9 | 21.8 | 21.8 | 22.0 | 22.1 | 21.8 | 20.6 | 21.8 | 1.4 | 4.2 | 0.0 |  |  | Cisse et al.^(^*^10^*^)^ | This study |
| P. murina (B123) | 22.4 | 22.3 | 22.4 | 22.5 | 22.6 | 22.3 | 21.0 | 22.3 | 12.3 | 14.7 | 12.5 | 0.0 |  | Ma et al. ^(^*^25^*^)^ | GCA_000349005.2 |
| P. wakefieldiae (2A) | 22.9 | 22.8 | 22.8 | 23.0 | 23.1 | 22.8 | 21.2 | 22.8 | 14.9 | 16.5 | 14.9 | 12.0 | 0.0 | This study | PRJNA632570 |

**Supplementary Table 6.** Subtelomeres in *P. macacae* (Excel spreadsheet)*.*

**Supplementary Table 7.** *P. jirovecii* genome-wide signatures of selection (Excel spreadsheet).
